# Evolutionary fingerprinting of protein-coding genes in RNA viruses

**DOI:** 10.64898/2025.12.05.692645

**Authors:** Laura Muñoz-Baena, Hugo G. Castelán-Sánchez, Sareh Bagherichimeh, Paula Magbor, Jorge Rojas-Vargas, Amjad Khan, Abayomi Olabode, Art F. Y. Poon

## Abstract

RNA viruses evolve rapidly to adapt to changing host environments. Much of this adaptation occurs at the level of protein-coding genes. Thus, some of the most well-characterized examples of rapid adaptation have been found in virus proteins that are exposed on the surface of the viral particle, where they mediate host receptor binding and cell entry. To investigate whether surface-exposed proteins and other proteins encoded by viruses exhibit different patterns of evolution under selection, we analyzed 244 protein-coding genes from 28 species of RNA viruses representing 15 taxonomic families. First, we show that gene-wide rates of non-synonymous (*dN*) and synonymous (*dS*) substitutions do not differentiate between categories of proteins. To provide a more detailed comparison between genes, we inferred for each alignment the bivariate posterior distribution over a fixed grid of codon site-specific *dN* and *dS* values. This distribution is the gene’s ‘evolutionary fingerprint’. Next, we computed the Wasserstein distance for every pair of fingerprints, which is analogous to amount of work required to reshape one distribution to another. After compensating for differences in genetic variation among viruses and proteins, we found that surface-exposed proteins could not be distinguished from non-exposed proteins in the space induced by the Wasserstein distance matrix. However, surface-exposed proteins from enveloped viruses were significantly clustered apart from their counterparts in non-enveloped viruses. In contrast, there was no significant separation between these categories of viruses for proteins with polymerase activity. We show that this pattern is more consistent with relaxed purifying selection than adaptive evolution in proteins associated with viral envelopes.

## Introduction

Viruses have remarkably high rates of molecular evolution [1]. In particular, elevated mutation rates in RNA viruses, attributed to the low replication fidelity of the virus-encoded RNA-dependent RNA polymerase [2], can provide an abundance of raw material for a rapid response to selection. Selection in virus populations is predominantly shaped by their host environments. This environment may include the host cell receptor proteins targeted by a virus for binding [3], cellular components that are incorporated into the virus replication cycle [4], and both innate and adaptive immune responses [5]. Of these potential factors, the host adaptive immune response is arguably the most diverse and capable of changing at the compressed time scale of RNA virus evolution. Much of what we understand about selection in viruses comes from protein-coding genes [6]. In general, the proportion of RNA virus genomes that encodes proteins (*i.e.*, the coding density) is relatively high. The proteins encoded by a virus genome can be broadly categorized into structural and non-structural proteins, depending on whether the protein becomes part of the viral particle or remains within the cell. Some structural proteins comprise the outer capsid of non-enveloped viruses, or become embedded in the membrane of enveloped viruses. These surface-exposed proteins are the primary interface between the virus and the extracellular host environment. For instance, surface envelope glycoproteins such as HIV-1 gp120 [7] and influenza A virus hemagglutinin [8] are well-characterized targets of selection by neutralizing antibodies. Consequently, comparative studies of selection in viruses have tended to focus on the surface-exposed proteins, *e.g.*, [9, 10]. On the other hand, significant positive selection has also been reported for genes encoding non-structural proteins or structural proteins that are not exposed on the surface of the virus particle [11].

This study endeavours to identify general trends in the types of selection experienced by different types of virus proteins, with a focus on testing the hypothesis that surface-exposed proteins undergo more positive selection than other viral proteins. Selection in protein-coding genes is typically identified by comparing the rates of amino acid-replacing (non-synonymous) and silent (synonymous) substitutions. When adjusted for the expected numbers of non-synonymous and synonymous substitutions, these rates become normalized quantities denoted respectively as *dN* and *dS* [12]. A relative excess of non-synonymous substitutions (*dN > dS*) provides evidence of positive selection, where selection promotes amino acid changes. In a constant and uniform selective environment, positive directional selection is a transient phenomenon that is resolved when a beneficial mutation becomes fixed in the population [13]. If we follow a single virus lineage through different host environments over time, the varying immune responses may lead to an excess of non-synonymous within-host polymorphisms [14]. Kistler and Bedford [15] recently demonstrated that lagging partial herd immunity can drive a sustained excess of non-synonymous substitutions in surface-exposed virus proteins. However, their analysis was limited to viruses with longitudinal samples of infections related by a single trunk lineage, *i.e.*, a ladder-like tree, because it relied on a comparison between substitutions and polymorphisms within a lineage (and its transient descendants) over time [16].

A more conventional approach to measuring selection in protein-coding genes in viruses is to fit a codon-substitution model across many divergent lineages that descend from a common ancestor [17]. In this context, positive selection driven by variation in selective environments is known as diversifying selection. This approach enables us to evaluate a broader selection of viruses. In this study, we examine whether different categories of virus proteins, including surface-exposed and non-exposed proteins, experience significantly different types of selection. Both synonymous and non-synonymous substitution rates can vary substantially among codon sites in a gene sequence

[18]. Estimating these codon site-specific rates with reasonable accuracy requires a substantial amount of genetic variation in the sequence alignment. As a result, we focused specifically on a curated set of twenty-eight RNA virus species with a sufficient number of publicly available full-length genome sequences with an adequate level of evolutionary divergence. We first demonstrate that standard methods that reduce each alignment to a summary statistic, *e.g.*, the gene-wide dN/dS ratio, do not resolve significant differences between exposed and non-exposed proteins. This implies that we require a more detailed method to compare site-specific patterns of selection between genes. The primary obstacle to this approach is that it is not obvious how one should compare site-level quantities between genes with no homology; for instance, HIV-1 envelope glycoprotein gp120 and enterovirus A71 helicase.

Pond and colleagues [19] described a statistical method to overcome this problem, which they dubbed ‘evolutionary fingerprinting’. The basic premise is that the rate variation among sites for a protein-coding gene alignment can be modeled as a latent discrete bivariate probability distribution over an *a priori* fixed grid of *dN* and *dS* rates. A flat prior distribution over this grid is updated by the phylogenetic likelihood of the codon alignment. The resulting posterior distribution over the grid is the evolutionary fingerprint of that alignment. Hence, the evolutionary fingerprint provides a common framework for comparing unrelated genes. We identify significant challenges that arise in applying fingerprinting to alignments from a broad diversity of rapidly-evolving species and genes, and develop methods to address these issues that have not been described in previous work [*e.g.*, 20–22]. Using this approach, we determine that there is not significant evidence that surface-exposed proteins undergo selection any differently than any other virus proteins. However, we observe that surface-exposed proteins associated with enveloped viruses have evolutionary fingerprints that are readily distinguishable from the fingerprints of their counterparts in non-enveloped viruses. Finally, we assess the implications of these results in light of what is currently known about the structure and function of these proteins.

## Methods

### Data collection

We manually queried the NCBI Genbank database to select candidate RNA virus species on the basis of two criteria: the availability of at least 100 complete or near-complete genome sequences, and the existence of an annotated reference genome, *i.e.*, RefSeq record [23]. In addition, we excluded sequences associated with patents, laboratory clones or modified nucleic acids. For each candidate species, we filtered the search results by taxonomic identifier and minimum sequence length based on the expected genome length, and then downloaded a list of accession numbers. If the virus had a segmented genome, *e.g.*, influenza A virus, then we manually composed queries including gene identifiers and exported separate lists of accession numbers. We anticipated that the genetic diversity captured in human immunodeficiency virus type 1 (HIV-1) sequences would be disproportionately greater than other viruses due to extensive sequencing of HIV-1 infections at a global scale. Consequently, we restricted our analysis to infections classified as sub-subtype A1, which is predominantly found in east Africa and central Asia [24]. We queried the Los Alamos National Laboratory HIV Sequence Database (https://www.hiv.lanl.gov) for sub-subtype A1, limiting the search results to one record per individual, and then extracted the Genbank accession numbers for subsequent steps. For influenza A virus (IAV), sequences from human hosts (predominantly subtypes H3N2 and H1N1) tend to induce highly ladder-like trees due to short infectious periods and transient cross-immunity in the host population [25]. This scenario is not consistent with diversifying selection as measured by comparative dN/dS methods [13]. Consequently, we restricted our search to subtype H9N2 infections isolated from avian hosts, where multiple co-circulating lineages with low pathogenicity have become endemic in commercial poultry [26].

For virus genomes in which genes were annotated separately as ‘mat peptide’ features, we used the BioPython [27] interface to the NCBI Entrez API to retrieve all protein-coding sequences (CDSs) associated with the records corresponding to a given set of accession numbers. This yielded a FASTA file containing multiple sequence records for every genome, with sequence labeled with protein name, genome strand and genome coordinates. We used the same script to extract sample metadata from the SeqRecord object, *e.g.*, sample collection date. Next, we used MAFFT (version 7.49) [28] to align the amino acid translation of each sequence against the set of mature peptides from the reference genome. We assigned each sequence to the reference peptide that attained the highest alignment score, given a match score of +1, a mismatch penalty of −1 and a linear gap penalty of −3. For virus genomes in which proteins are derived from a polyprotein encoded by a single open reading frame, *e.g.*, hepatitis C virus, we used a similar pairwise method to align the polyprotein sequence pairwise to every mature peptide feature in the reference genome, and extracted the corresponding nucleotide substring to a separate FASTA file for each feature.

We classified each protein-coding gene as ‘surface-exposed’ if any portion of the mature peptide was documented (*e.g.*, ViralZone [29]) to be exposed on the outer surface of the virus particle released into the extracellular environment. Proteins from plant viruses were not labeled as ‘surface-exposed’ even if they are exposed on the surface of the virus particle, because host plants do not have an adaptive immune system. In addition, we annotated genes that encode enyzmatic proteins with polymerase or protease activity, or structural proteins. We annotated virus species by whether they are enveloped or non-enveloped.

### Phylogenetic analysis

For each FASTA file produced in the preceding step, we used a Python script to generate a multiple alignment of the amino acid translations of the sequences using MAFFT, and then applied the gaps in this alignment to the original nucleotide sequences to obtain a codon alignment preserving the reading frame. We used AliView [30] to visually inspect the resulting alignment, and manually removed problematic sequences, *e.g.*, CDS records labeled with the wrong protein. Positions where a majority of sequences contained a gap were removed from the alignment in a codon-aware manner. Incomplete sequences that were shorter than half of the alignment length were excluded. We used FastTree (version 2.1.11, compiled for double precision) [31] to reconstruct a preliminary maximum likelihood tree from the resulting alignment. The tree was visually inspected for excessively long terminal branches. Any outlier sequences identified at this step were removed and the tree was rebuilt from the updated alignment.

Our preliminary analyses indicated that measuring evolutionary fingerprints (*i.e.*, the posterior distribution of *dN* and *dS* values, see below) was sensitive to the amount of genetic variation captured by the alignment. We used the tree length, *i.e.*, sum of branch lengths to quantify this genetic variation. To determine the lowest acceptable tree length, we used INDELible version 1.03 [32] to simulate codon sequence alignments with known site-specific *dN* and *dS* values. We seeded the simulation with a random tree relating 100 tips that was generated under a constant size coalescent model. This input tree was rescaled to different lengths and the resulting alignments were analyzed using the FUBAR method (described in the next section). We calculated the root mean square error (RMSE) between the known and estimated *dN/dS* ratios. Based on our observation of a drop in RMSE associated with tree lengths greater than 1.0 expected substitutions per codon site and diminishing returns with increasing tree length (Supplementary Figure S1), we chose a target range of 0.5 to 2.0 expected nucleotide substitutions per site (ESS). To normalize tree lengths across alignments, we progressively removed the shortest terminal branches from the starting tree until the length fell below a cutoff of 2.0 ESS. This pruning approach maximized the amount of genetic variation for a given subset of sequences. In many cases, it was not possible to prune the tree down to the cutoff length because of long internal branches in the tree. For these alignments, we arbitrarily selected a starting terminal branch and proceeded toward the root until we reached an internal node that rooted a monophyletic clade with a total length below the cutoff. If the starting tree length was already between 0.5 and 2.0 ESS, then the alignment was passed to next steps without modification. Alignments with a starting tree length below 0.5 ESS were discarded from further analysis.

### Selection analysis

Many virus genomes contain overlapping genes in different reading frames as a potential mechanism for increasing the information content of a compact genome [33]. Overlaps of protein-coding genes cause problems for measuring selection because a substitution that is synonymous in one reading frame may be non-synonymous in another [34]. To reduce the influence of overlapping genes, we manually removed codon sites affected by overlaps in each alignment. Genes encoding multiple products due to alternate initiation or termination sites were not modified (*e.g.*, VP7 in rotavirus A), unless one of those alternate products involved splicing a frame-shifted portion of the gene (*e.g.*, influenza A virus PB2-S1). Any intervals involved in an overlap with different reading frames was removed from all affected gene alignments. Detailed reference coordinates of the final gene alignments are provided in Supplementary Table S1.

For each gene alignment, we reconstructed a maximum likelihood phylogeny using FastTree and then fit a Muse-Gaut codon substitution model crossed with a general time-reversible model of nucleotide substitutions in HyPhy to measure the gene-wide *dN/dS* ratio as a global parameter. We used the single likelihood ancestor counting (SLAC) method to estimate individual codon site-specific *dN* and *dS* values. Because gene-wide *dN/dS* ratios were right-skewed and strictly positive, we used a gamma regression model with a log-link function to evaluate the effects of whether the *i*-th virus is enveloped (*e_i_*) and the *j*-th protein of the virus is surface-exposed (*x_i j_*) on *dN/dS*. To account for variation in *dN/dS* = *ω* among virus species, we fit a mixed-effects log-link model using the R package *lme4* [35]:

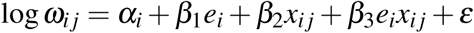

where *α_i_* is a random (species-specific) intercept term, *β*_•_ are fixed effects, and *ε* represents residual (error) variance. We generated 95% confidence intervals (CI) for model parameters by bootstrap resampling, and reported these intervals alongside the *P*-values. A term was considered to have a significant fixed or random effect if the 95% CI did not include zero. In addition, we used a binomial regression model with a logit link function to examine the association between these predictors and the proportion of codon sites under significant (*α* = 0.1) diversifying (*dN > dS*) selection:

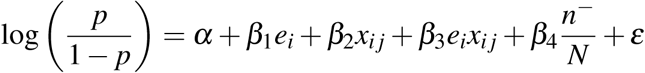

where *n*^+^ is the number of codon sites under significant diversifying selection, *n*^−^ is the number of sites under significant purifying selection, *N* is the total number of codon sites, and *p* = *n*^+^*/*(*N* − *n*^−^).

### Evolutionary fingerprinting

We used the Fast Unconstrained Bayesian Approximation (FUBAR) method [36] in *HyPhy* (version 2.5.60) [37] to estimate the site-specific synonymous (*dS*) and non-synonymous (*dN*) substitution rate parameters for each codon alignment. This method approximates a latent bivariate distribution of continuous *dN* and *dS* values with a discrete posterior distribution over a fixed 20 × 20 grid of values, which Pond *et al.* [19] dubbed the ‘evolutionary fingerprint’ for the alignment. Following Murrell *et al.* [21], we replaced the default grid values in FUBAR with a smoother distribution of rates generated by the formula (50 × *k*^5^)*/*19^5^ for *k* = {0, 1*,…,* 19}. In addition, we increased the length of the chain sample from the default 2 × 10^6^ to 10^7^ steps to improve sample convergence.

We used the Wasserstein distance (also known as the earth mover’s distance) implemented in the R package *transport* [38] to compare fingerprints obtained from two different alignments. This distance is analogous to the minimum amount of work required to reshape one distribution to another, accounting for the distance between points on the grid. We used the Euclidean norm for calculating this distance. The resulting distance matrix was visualized by multidimensional scaling using the R function *cmdscale*. Using this analysis, we determined that evolutionary fingerprints were also sensitive to alignment length. Intuitively, the number of codon sites in the alignment roughly corresponds to sample size. To address this effect for alignments longer than a given threshold, *e.g.*, *L* = 50 codon sites, we generated 10 replicate subsets by sampling *L* codon sites from the alignment at random without replacement. We processed the sample alignments using the same workflow, and then calculated the centroid for each set of replicates by averaging their coordinates in the multidimensional scaling projection. This analysis was also repeated with a higher threshold of 100 codon sites.

### Supervised learning

We used a weighted *k*-nearest neighbour (*k*-NN) learning method to evaluate our ability to classify virus proteins by their evolutionary fingerprints. This supervised learning method simply assigns a label to each point in the test set by taking the majority consensus of its weighted *k* nearest neighbours in the training set [39]. The closest neighbors were obtained from the Wasserstein distance matrix. Because of the limited amount of data available, particularly when restricting the analysis to a specific category of proteins, we could not use a full train-test-validation framework to determine the optimal number of neighbors (*k*). Instead we obtained *a priori* results for *k* = 5 and then characterized the effect of varying *k* afterwards. In addition, we used a distance-weighted variant of the *k*-NN algorithm to account for the imbalanced frequencies of labels, where the contribution of each neighbour is weighted by its inverse distance from the query sample. The *j*-th query sample is assigned the label *y* that maximizes the following weighted sum:

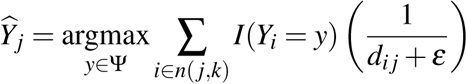

where Ψ is the set of all labels; *n*( *j, k*) is the set of the *k*-nearest neighbours of *j*; *I*(*x*) is an indicator function that assumes the value 1 if *x* is true and 0 otherwise; *d_i j_* is the Wasserstein distance from *i* to *j*; and *ε* is a small constant (in our case, 10^−9^) to make the algorithm numerically stable to identical samples [40]. We evaluated binary classification tasks for five sets of labels: (i) surface-exposed versus non-exposed proteins, (ii) polymerase vs. non-polymerase proteins, (iii) surface vs. polymerase, (iv) enveloped vs. non-enveloped viruses for only surface-exposed proteins, and (v) enveloped vs. non-enveloped viruses for only polymerase proteins.

All sequence data at different stages of processing have been deposited into a public online repository at https://doi.org/10.5281/zenodo.16320684 under a permissive license (Creative Commons Attribution 4.0 International). Python and R scripts implemented for this study have been published under the MIT license at https://github.com/PoonLab/surfaces.

## Results

### Gene-wide selection

We obtained protein-coding sequences from over 42,000 genomes for 28 different RNA virus species representing 15 different families (Table 1). These data largely comprised human viruses that pose a significant threat to human health, but we did not limit our analysis to viruses from human hosts — we also obtained data for several agriculturally-significant plant RNA viruses such as potato virus Y [41]. Figure 1A displays the mean estimates of dN/dS for each alignment of protein-coding gene sequences. As expected, all mean dN/dS values were well below 1, indicating that a majority of codon sites in any given gene were under purifying selection. For some viruses such as HIV-1, proteins that are located on the surface of the virus particle and exposed to an adaptive host immune response tended to have a higher mean dN/dS (*e.g.*, gp120; Figure 1). Hence, we evaluated support for the hypothesis that surface-exposed virus proteins have relatively more sites under diversifying selection, which would drive up the mean dN/dS ratio. We note that surface-exposed proteins for viruses infecting plants were excluded from this category because plant hosts do not have an adaptive immune response that would drive diversifying selection. Ignoring variation in mean dN/dS values among viruses, we found no significant effect of surface exposure in a gamma regression model (*P* = 0.09, 95% CI = −0.03, 0.56), where we used a log-link function to account for the skew of this ratio outcome. If we switched to a mixed-effects log-link model to address the inherent structure in these data (*i.e.*, repeated measures from each virus species), we observed significant variation among viruses (*P <* 10^−6^, standard deviation 95% CI = 0.68, 1.12) and a marginally significant effect of surface exposure (*P* = 0.04, 95% CI = −0.05, 0.37). Adding an interaction between surface exposure and being an enveloped virus conferred an improved model fit (ΔAIC = 6.7), but the main effect of surface exposure (*P* = 0.48, 95% CI = −0.47, 0.23) becomes absorbed into the interaction term (*P* = 0.03, 95% CI = −0.05, 0.85). These trends are broadly consistent with, but do not directly support, the hypothesis that surface-exposed proteins tend to experience a different selective regimen than other virus proteins.

**Figure 1:**
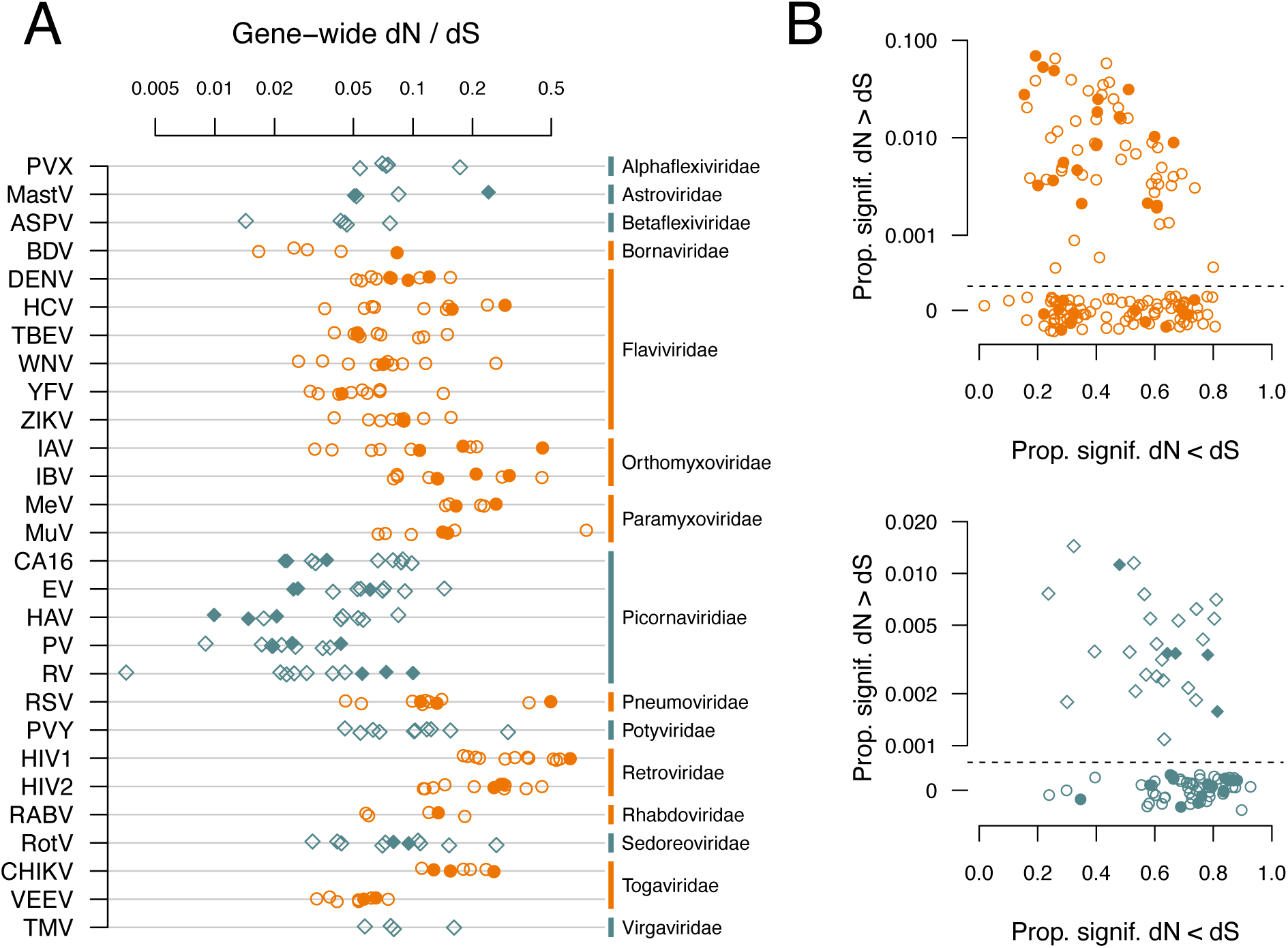
Summary of dN and dS estimates for protein-coding gene alignments. (A) Gene-wide dN/dS estimates per protein and virus. Each point represents a gene alignment, grouped by virus (left labels) and by family (right labels). A point is filled if the protein is located on the surface of the virus particle and potentially exposed to an adaptive immune response. Colour and shape are used to distinguish proteins from enveloped (circle, orange) and non-enveloped (diamond, blue) viruses. (B) Scatterplots of the proportions of codon sites under statistically significant (*α* = 0.1) purifying (*dN < dS*, *x*-axis) and diversifying (*dN > dS*, *y*-axis) selection for enveloped (top) and non-enveloped (bottom) viruses. Each point represents a gene alignment, using the same shape and colour scheme as (A). The *y*-axis was log-transformed to accommodate skewed distributions in the proportions of *dN > dS* sites, with an axis break for zero counts, including random noise to reduce overlap.

**Table 1:** Summary of viruses analyzed in this study. *Abbrv.* = conventional abbreviation used for figures. *Env?* = is enveloped virus? *Genomes* = initial number of genome records obtained prior to filtering. *For viruses with segmented genomes, we reported the maximum number of sequences for any segment. *Proteins* = number of protein-coding genes analyzed for selection; note this number excludes genes of insufficient length or with extensive overlaps with other genes.

| Family | Virus | Abbrev. | Env? | Genomes | Proteins |
| --- | --- | --- | --- | --- | --- |
| Alphaflexiviridae | potato virus X | PVX | No | 409 | 5 |
| Astroviridae | mamastrovirus | MastV | No | 125 | 4 |
| Betaflexiviridae | apple stem pitting virus | ASPV | No | 166 | 5 |
| Bornaviridae | orthobornavirus | BDV | Yes | 118 | 5 |
| Flaviviridae | dengue virus 2 | DENV | Yes | 1,630 | 10 |
|  | hepatitis C virus (genotype 1a) | HCV | Yes | 3,911 | 10 |
|  | tick-borne encephalitis virus | TBEV | Yes | 440 | 11 |
|  | West Nile virus | WNV | Yes | 2,313 | 11 |
|  | Yellow fever virus | YFV | Yes | 1,304 | 10 |
|  | Zika virus | ZIKV | Yes | 664 | 10 |
| Orthomyxoviridae | influenza A virus (H9N2) | IAV | Yes | 2,220* | 10 |
|  | influenza B virus | IBV | Yes | 13,224* | 10 |
| Paramyxoviridae | measles virus | MeV | Yes | 825 | 6 |
|  | mumps virus | MuV | Yes | 721 | 7 |
| Picornaviridae | coxsackievirus A16 | CA16 | No | 514 | 10 |
|  | enterovirus A71 | EV | No | 1261 | 10 |
|  | hepatitis A virus | HAV | No | 362 | 9 |
|  | polio virus | PV | No | 270 | 10 |
|  | rhinovirus A | RV | No | 1347 | 10 |
| Pneumoviridae | respiratory syncytial virus | RSV | Yes | 1,021 | 11 |
| Potyviridae | potato virus Y | PVY | No | 911 | 10 |
| Retroviridae | human immunodeficiency virus type 1 (subtype A) | HIV1 | Yes | 256 | 12 |
|  | human immunodeficiency virus type 2 | HIV2 | Yes | 121 | 12 |
| Rhabdoviridae | rabies virus | RABV | Yes | 2,229 | 5 |
| Sedoreoviridae | rotavirus A | RotV | No | 4,635* | 11 |
| Togaviridae | Chikungunya virus | CHIKV | Yes | 978 | 7 |
|  | Venezuelan equine encephalitis virus | VEEV | Yes | 220 | 9 |
| Virgaviridae | tobacco mosaic virus | TMV | No | 108 | 4 |

Measuring selection at the level of whole genes may obscure more significant effects on a small number of codon sites. Typically only a fraction of the amino acids in a protein are actually exposed on the surface of the virus, for instance [7]. Figure 1B summarizes the proportions of codon sites under significant site-specific diversifying (*dN > dS*) or purifying (*dN < dS*) selection for each gene. We fit a regression model to the number of sites with significant *dN > dS* as a binomial outcome accounting for the total number of sites, with the proportion of remaining sites with significant *dN < dS* as an independent variable, and with surface-exposure and enveloped virus as additional main and interaction effects. Models dropping any of these independent variables were rejected (ΔAIC ≥ 3.9). The number of sites with diversifying selection declined significantly with the amount of purifying selection at other sites (*P* = 0.016, 95% CI = −1.51, −0.16). There was no significant association with surface exposure (*P* = 0.57, 95% CI = −1.12, 0.51) or enveloped virus (*P* = 0.35, 95% CI = −0.18, 0.55), but the interaction effect departed significantly from zero (*P* = 0.025, 95% CI = 0.18, 1.89).

### Evolutionary fingerprints

Ideally, we want to compare sets of site-specific estimates of *dN* and *dS* between two genes, rather than comparing a single number, such as the gene-wide average *dN/dS* or proportion of sites with significant diversifying selection. The genes will usually be non-homologous and can differ substantially in length. Consequently, it is not feasible to align codon sites from different genes in a meaningful way. Pond *et al.* [19] proposed a method to characterize a gene by assuming that the site-specific rates are drawn from a latent bivariate probability distribution of *dN* and *dS*, dubbed the ‘evolutionary fingerprint’. By constraining this distribution over a fixed grid of *dN* and *dS* values, this fingerprint provides a common framework in which one can compare completely unrelated genes. We employed this method to generate the fingerprints for the protein-coding gene alignments in our study. However, we found that the shape of an evolutionary fingerprint is sensitive to the number or diversity of sequences in the alignment. Specifically, longer trees corresponding to larger and/or more diverse sequence alignments tend to yield more granular fingerprints for the same protein (Supplementary Figure S2). To control for this effect, we constrained the tree lengths to fall within the range of 0.5 to 2.0 expected nucleotide substitutions per site (Supplementary Figure S3). In addition, we determined that the length of the alignment, *i.e.*, the number of codon sites (*L*), had a substantial effect on evolutionary fingerprints (Supplementary Figure S4), which we addressed by sampling *L* = 50 or *L* = 100 sites at random without replacement from each codon alignment. A random selection of fingerprints for *L* = 100 are displayed in Figure 2. Generally, the posterior probability mass is concentrated in the middle of the range of *dS* values, and largely below the *dN* = *dS* line of neutral evolution, *i.e.*, in the purifying selection regime. While this visualization reveals substantial variation in fingerprints among genes, it is difficult to use these to discern patterns with the unassisted eye.

**Figure 2:**
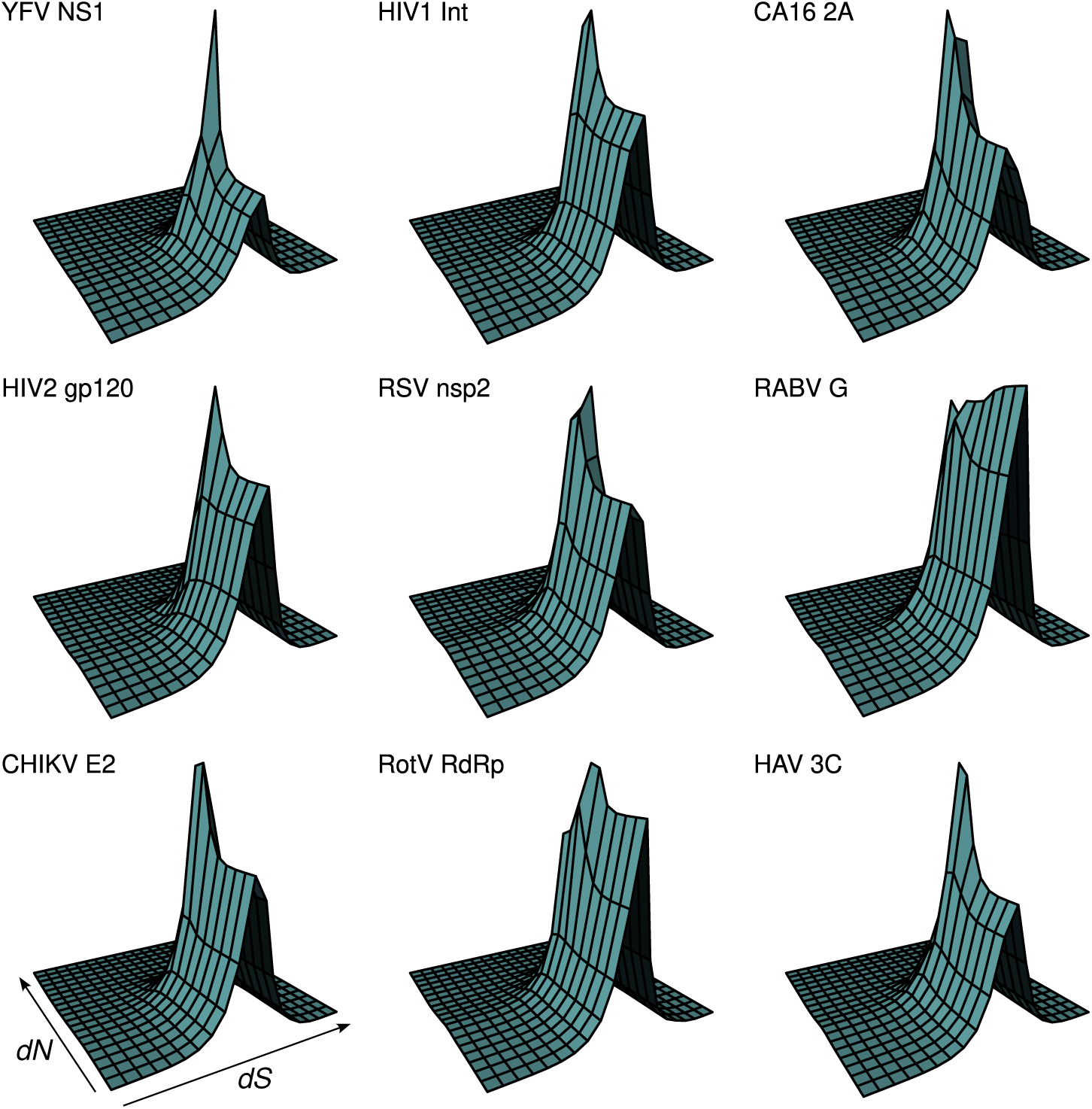
Evolutionary fingerprints for a random sample of *n* = 9 virus proteins. An evolutionary fingerprint is the estimated posterior probability distribution on a fixed 20 × 20 grid of *dN* (*y*-axis) and *dS* (*x*-axis) values. Each perspective plot depicts the fingerprint averaged over 10 random samples of 100 codon sites from a gene alignment, labeled with the abbreviated virus (see Table 1) and protein names.

To analyze the evolutionary fingerprints in a quantitative framework, we calculated the Wasser-stein distance between every pair of fingerprints. This distance roughly corresponds to the amount of ‘work’ required to transform one distribution into another. Figure 3 displays multidimensional scaling (MDS) plots that reduce the resulting pairwise distance matrix into two dimensions. Each point represents the average (centroid) of ten replicate samples of 100 codon sites from a given gene alignment. Replicate samples formed distinct clusters when visualizing the entire distance matrix (Supplementary Figure S5). Although this sampling procedure substantially reduced the effect of variation in alignment lengths (Supplementary Figure S4), requiring a minimum of 100 codons excluded 41 (16.8%) out of 244 gene alignments from our analysis; the median alignment length was 255.5 (interquartile range, IQR: 130 − 466 codons; Supplementary Figure S3). Consequently, we repeated our analysis using a smaller sample of 50 codon sites per alignment and obtained similar results (Supplementary Figure S6).

**Figure 3:**
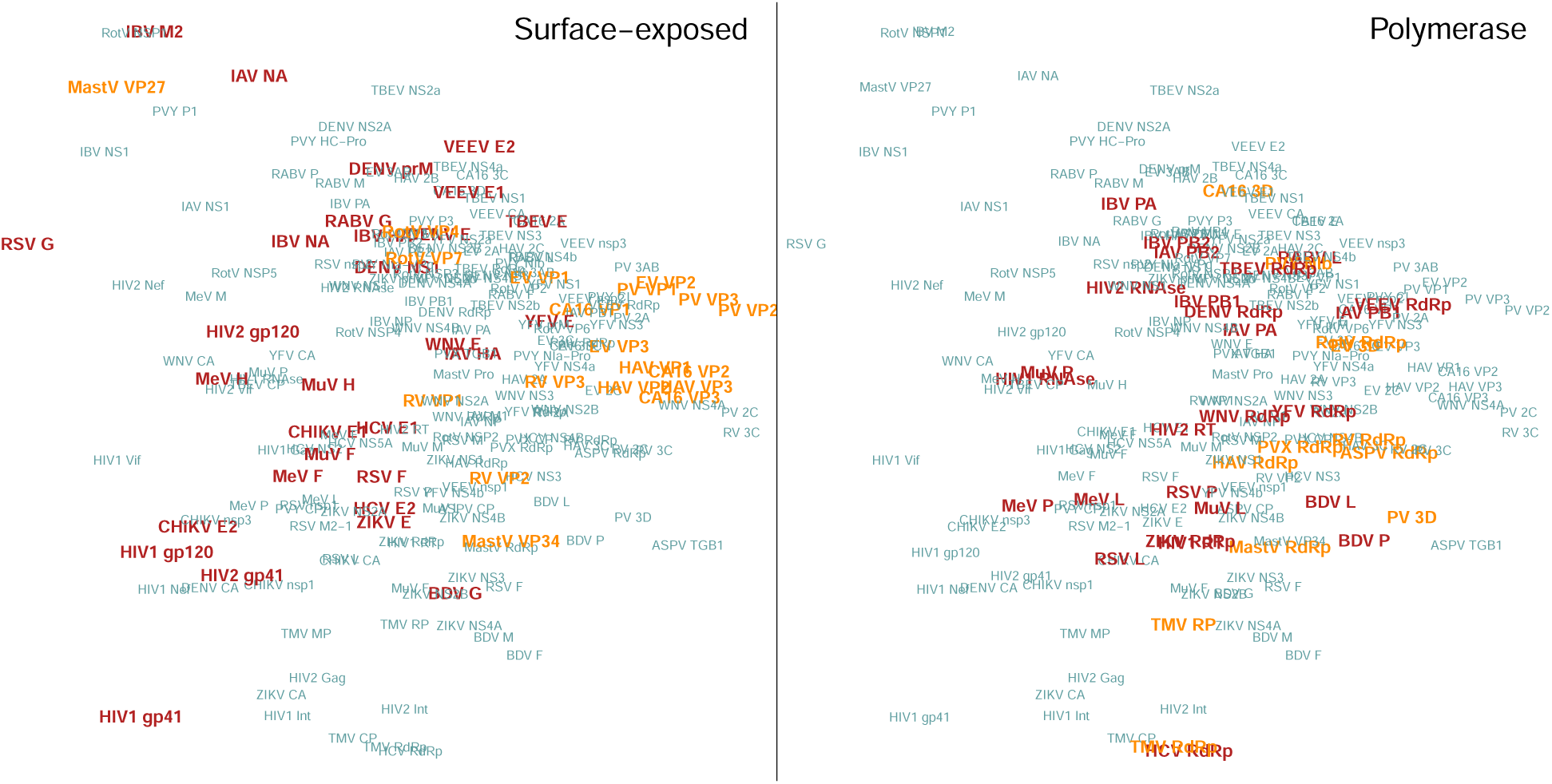
Multidimensional scaling plots of the Wasserstein distance matrix. Each point represents the centroid of ten replicate samples of *L* = 100 codon sites from a gene alignment, labeled with the respective virus and protein (abbreviations defined in Table 1 and Supplementary Table S1). The *x*− and *y*-axes capture 51% and 15% of the variance, respectively. Labels in the left plot are drawn at a larger size if the protein was classified as surface-exposed, and shaded to indicate that the exposed protein is carried by an enveloped (darker, red) or non-enveloped (lighter, yellow) virus. In the right plot, labels are larger for proteins with polymerase activity, with the same darker shading for enveloped viruses.

Overall, we did not see any clear separation between surface-exposed and non-exposed proteins in this space. However, if we focused on the surface-exposed proteins (*n* = 49), we found that they separated with respect to their association with enveloped and non-enveloped viruses, respectively (Figure 3, left). In particular, the VP1, VP2 and VP3 proteins that make up the outer surface of the capsid of enteroviruses (*i.e.*, coxsackievirus A16, enterovirus A71, hepatitis A virus, poliovirus, rhinovirus) formed a distinct cluster. For contrast, we generated a similar plot high-lighting proteins with polymerase activity, which affected a similar number of alignments (*n* = 38; Figure 3, right). These enzymatic proteins are generally not surface-exposed, although some are technically structural because they become incorporated into the virus particle (*e.g.*, HIV-1 reverse transcriptase). The corresponding fingerprints tend to map to a different region of the MDS projection compared to the surface-exposed proteins. We observed that there was less of a discernible separation between polymerases from enveloped and non-enveloped viruses.

### Supervised learning

The MDS visualization in Figure 3 implies that surface-exposed proteins do not share a characteristic evolutionary fingerprint that is distinct from proteins that are not exposed. On the other hand, the same visualization suggests that the surface-exposed proteins associated with non-enveloped viruses can be separated from those associated with enveloped viruses. To test this observation, we applied the *k*-nearest neighbours (*k*-NN) method to the Wasserstein distance matrix to classify fingerprints with respect to five sets of labels, including the above comparison (restricted to the subset of surface-exposed proteins). Due to the limited sample size (*i.e.*, *n* = 203 alignments for samples of *L* = 100 codon sites, or *n* = 49 surface-exposed proteins), we used leave-one-out cross-validation to assess classifier performance. Results for surface proteins associated with enveloped or non-enveloped viruses (*L* = 100) are summarized in Figure 4. While a majority of points were assigned the correct labels using *k*-NN, there were several points in the overlapping region between clusters that were misclassified. In the replicate depicted in Figure 4A, for example, there were five misclassified surface-exposed proteins from enveloped viruses, specifically influenza A virus hemagglutinin (HA), orthobornavirus glycoprotein (G), tick-borne encephalitis virus envelope (E), Venezuelan equine encephalitis virus envelope (E2), and Yellow fever virus envelope (E). For non-enveloped viruses, the misclassified proteins were coxsackievirus A16 VP1, enterovirus VP1, mamastrovirus VP27, rhinovirus A VP1, and rotavirus A VP4 and VP7.

**Figure 4:**
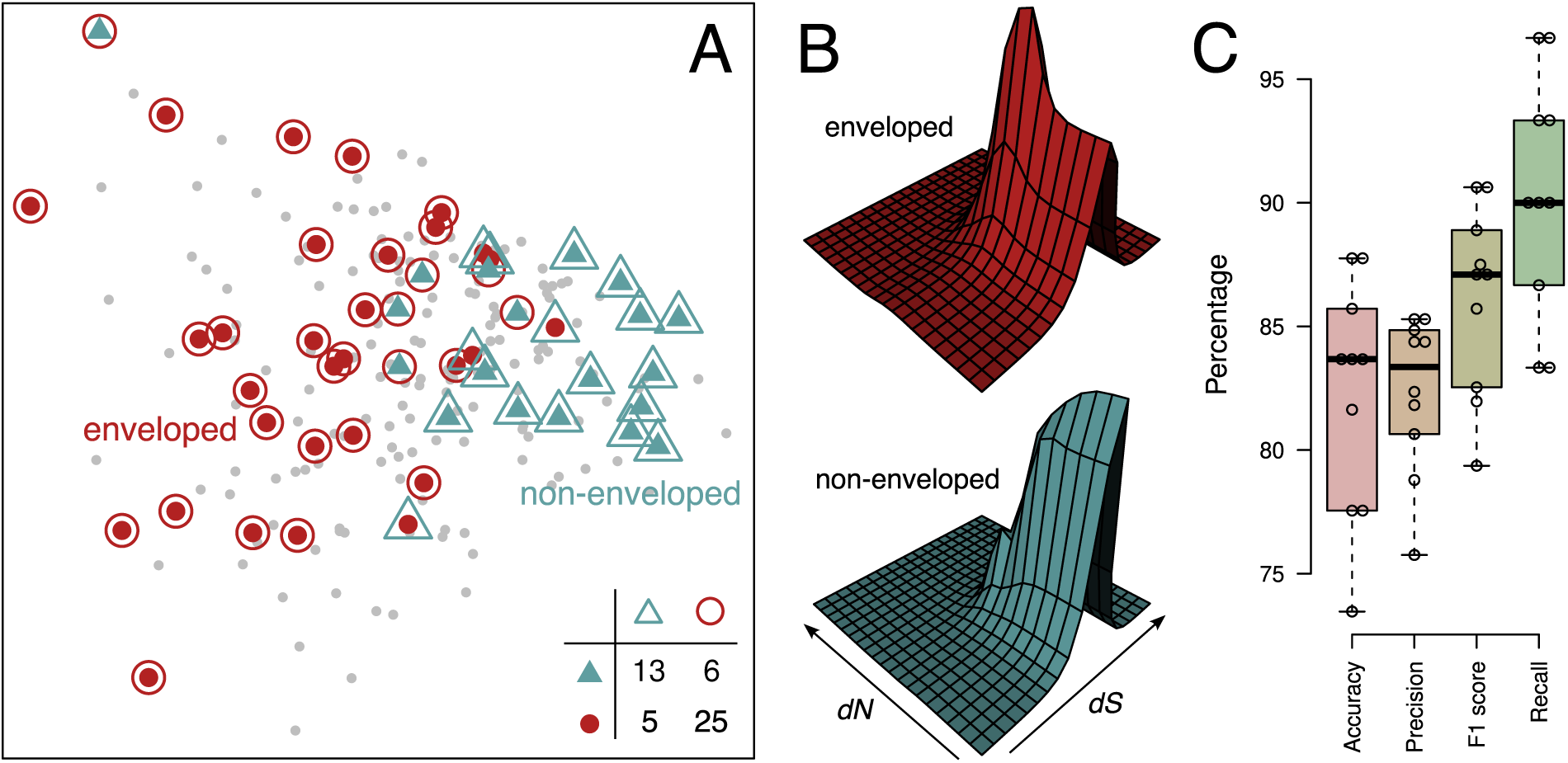
Summary of *k*-nearest neighbours (*k*-NN) analysis of evolutionary fingerprints for surface-exposed proteins. (A) This scatterplot depicts the multidimensional scaling projection of the Wasserstein distance matrix for the first of ten replicate, sampling *L* = 100 codon sites from *n* = 203 alignments. Each large point represents a surface-exposed protein from an enveloped (solid circle, red) or non-enveloped (solid triangle, blue) virus. Non-exposed proteins are represented by small, grey points. Open circles and triangles represent the predicted labels for *k* = 5; thus, discordant shapes indicate misclassified points. The confusion matrix is provided in the lower-right corner. (B) Perspective plots of the evolutionary fingerprints for surface-exposed proteins, averaged over *n* = 30 enveloped viruses (top) or *n* = 19 non-enveloped viruses (bottom). (C) Box-and-whiskers plots summarizing the performance of *k*-NN for *n* = 10 replicate sets of fingerprints for all ten replicates, using four different metrics. Detailed results are also provided as Supplementary Figure S7.

To visually assess the differences between these two groups of surface-exposed proteins, we averaged the evolutionary fingerprints across proteins for *n* = 30 enveloped and *n* = 19 non-enveloped viruses. We observed that the averaged distribution was shifted more towards the purifying selection regime for non-enveloped viruses (Figure 4B). This shift was also visualized by taking the difference between these distributions (Supplementary Figure S8). The heatmap not only indicates the region of *dN < dS* where evolutionary fingerprints for non-enveloped viruses are concentrated, but also revealed a cluster of elevated probabilities for intermediate *dN* and *dS* values in enveloped viruses.

Based on several performance metrics (Table 2), the *k*-NN method was the most effective at distinguishing between surface-exposed proteins from enveloped and non-enveloped viruses, relative to the other classification tasks. The second-best task was distinguishing between polymerases from enveloped and non-enveloped viruses, although the accuracy was poor. Higher accuracy values for comparisons involving the full dataset (*e.g.*, surface vs. non-exposed) relative to other performance metrics were most likely caused by class imbalance. We tended to obtain slightly better performance when we reduced the alignment length from *L* = 100 to 50 codons, which restored an additional 41 fingerprints to the training set.

**Table 2:**
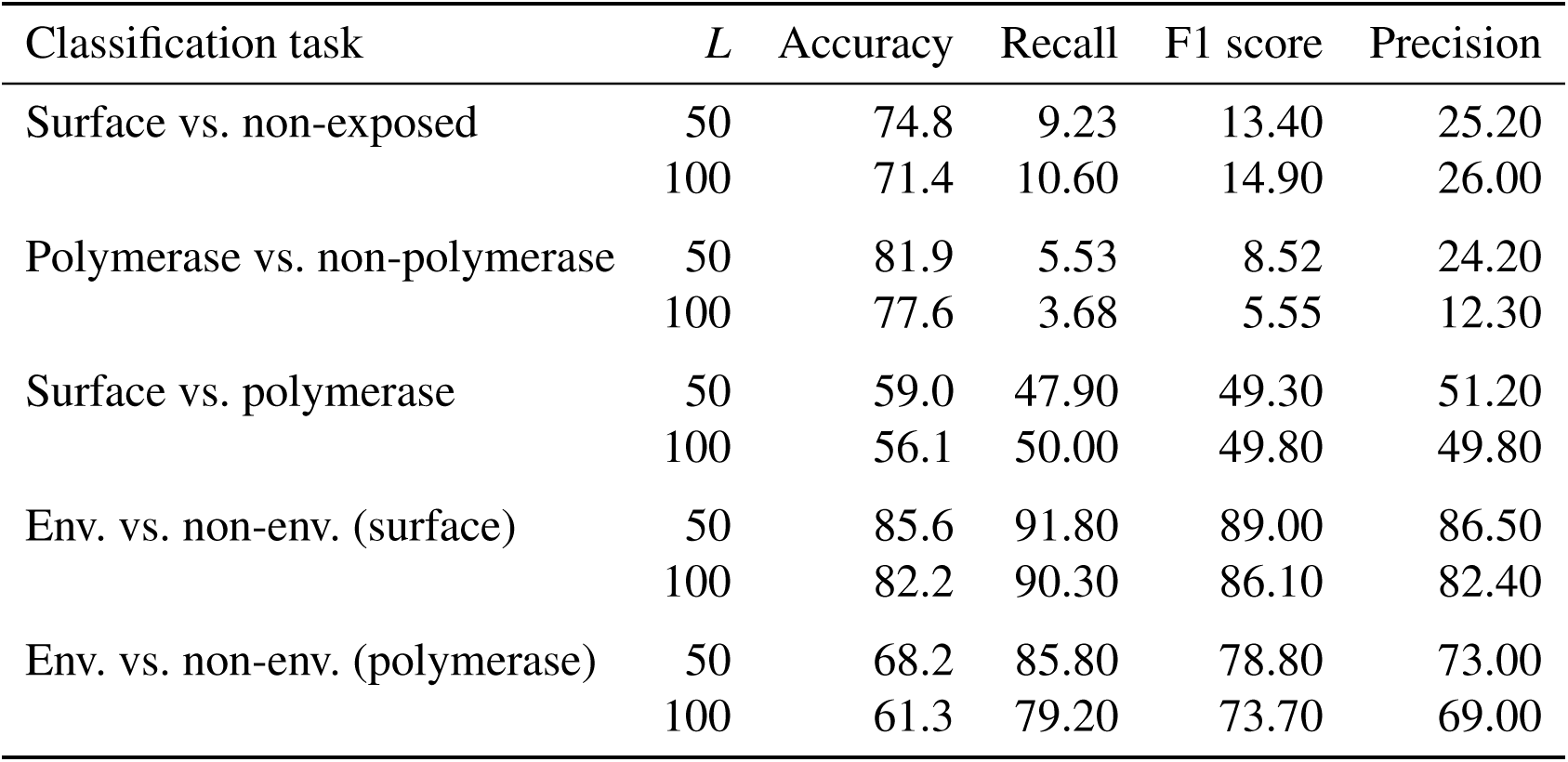
Classification accuracy, recall, F1 score, and precision (%) averaged over 10 replicates (*k* = 5) for *L* = 100 and *L* = 50, across five different clasification tasks. Higher is better performance.

## Discussion

Our results indicate that selection on surface-exposed proteins is not particularly different from other virus proteins, at least when measuring diversifying versus purifying selection on multiple co-circulating lineages in a host population. Hence, our analysis does not support the hypothesis that surface-exposed proteins are evolutionarily distinct from other proteins encoded by virus genomes due to selection by neutralizing antibodies. Although the humoral immune response is a major cause of selection in the host environment, there are many other factors that also contribute to virus evolution. The cellular immune response, for instance, can act on any protein produced from the virus genome, and is also an important part of the adaptive immune response. For example, an analysis of a large longitudinal dataset of HIV-1 genome sequences within a single subject [42] found that about half of codon sites under directional or diversifying selection were associated with genes other than *env*, and about 40% of these sites were associated with cytotoxic T-cell lymphocyte epitopes.

We also observed, however, that among surface-exposed proteins, those from enveloped viruses are affected by selection differently from their counterparts in non-enveloped viruses. This pattern was resolved the most clearly by evolutionary fingerprinting, after making adjustments for differences in genetic variation. Why might association with a viral membrane change the selective environment of a protein? We noted that the shift between mean fingerprints is consistent with the relaxation of purifying selection in enveloped viruses (Figure 4 and Supplementary Figure S8). Thus, a possible explanation is that maintaining the structural and functional integrity of a protein capsid may involve a greater number of conserved protein-protein interactions than membrane-embedded proteins. For example, a recent study [43] found that naturally-occurring viral capsids have more protein-protein interactions than synthetically-engineered capsids, or capsid-like structures produced by the overexpression of a structural protein. The scale of this effect may mask positive selection targeting a much smaller number of antigenic sites in surface-exposed proteins [8]. A caveat is that this separation between enveloped and non-enveloped viruses was not entirely exclusive to surface-exposed proteins; we observed some separation for proteins with polymerase activity as well. Even though classifier performance was inferior for polymerases, it remains possible that some of the separation in surface-exposed proteins may be confounded by generalized differences between these two groups of viruses.

A prominent outlier in our analysis of evolutionary fingerprints for surface-exposed virus proteins is the VP27 protein of the non-enveloped mamastrovirus. VP27 forms a dimer with VP25, a shorter gene product derived from the same precursor polypeptide following proteolytic cleavage by trypsin [44]. This precursor is encoded by the hypervariable central region of the genome [45]. The resulting dimer forms a spike that protrudes from the core structure of the virus particle and is highly antigenic [46]. Notably, the only other non-enveloped virus that encodes a spike protein in our collection was rotavirus A (VP4) [47] — the Picornaviridae have a spherical capsid with no distinct spike structures [48], and the plant viruses have a filamentous or rod-like structure formed by multi-functional coat proteins [49]. Both rotavirus VP4 and VP7, a glycoprotein forming up the outer surface of the capsid, induce neutralizing antibodies and are used to define serotypes. Like mamastrovirus VP27, both rotavirus proteins tended to be misclassified as coming from an enveloped virus (Figure 4).

Another interesting result from our analysis of evolutionary fingerprints is that VP1 proteins were substantially more likely to be misclassified as surface-exposed than the VP2 and VP3 major capsid proteins from the same enterovirus species, *i.e.*, rhinovirus, enterovirus A71, polio virus, and coxsackievirus A16. These three major capsid proteins are structurally similar, sharing a common *β*-sandwich jelly roll fold, and contribute jointly to the formation of the outer capsid in equal numbers. However, VP1 forms most of the ‘canyon’ at the centre of the pentameric subunit that is responsible for host receptor-binding. Thus VP1 is highly exposed with variable loops that are important targets for neutralizing antibodies [50, 51]. In addition, it been found to cluster apart from VP2 and VP3 with respect to sequence composition or physicochemical properties [52, 53].

We also observed that capsid proteins from Flaviviruses (West Nile virus, Dengue virus, tick-borne encephalitis virus, yellow fever virus, Zika virus and hepatitis C virus) tended to be located apart from a cluster of the other proteins from the same virus species in the MDS projection of evolutionary fingerprints (Supplementary Figure S9). This pattern was the most striking for Dengue virus. These capsid proteins dimerize to form a nucleocapsid that encapsulates the viral RNA genome. Several studies have characterized both cellular and humoral responses to capsid proteins in Flaviviruses, *e.g.*, [54]. Because these proteins are not exposed on the surface of the virus protein, however, they are not expected to produce a neutralizing antibody response or interfere with host cell binding and entry [55]. Capsid proteins in flaviviruses are also multifunctional and interact with a number of host factors [56]; however, this characteristic is not unique to capsid proteins [57]. Consequently, it is not readily apparent why capsid proteins in flaviviruses appear to have a distinct evolutionary fingerprint.

### Comparison to previous work

Kistler and Bedford [15] recently reported significantly greater adaptive evolution — defined as an excess number of non-synonymous substitutions — in surface-exposed proteins than their non-exposed counterparts in a number of viruses. Specifically they found significant adaptive evolution in 10 out of 28 receptor-binding proteins, and in none of 27 polymerase proteins. Derived from the McDonald-Kreitman (MK) test by Bhatt and colleagues [16], this statistic compares the observed number of non-synonymous substitutions (replacements) to the expected number, which is derived from the observed number of synonymous substitutions and the ratio of non-synonymous and synonymous polymorphisms. Bhatt *et al.* analyzed a broader selection of 82 virus species (95 proteins), but there was no significant association of adaptive evolution (MK test *p <* 0.05*/*95) with surface exposure (Fisher’s exact test, odds ratio 1.1; 95% CI 0.46, 2.7), which is more consistent with our results. A major difference between these studies and the present work is that our method requires that co-circulating pathogens experience different selective host environments, *i.e.*, diversifying selection. The *dN/dS* method does not consistently detect uniform directional selection from longitudinal samples of a single lineage [13]. On the other hand, this approach has fewer restrictions on sample collection and tree shape (substantial numbers of sequences sampled over a decade or more, and a ladder-like phylogeny), making it amenable to apply to a broader range of viruses.

A substantial difference between [15] and [16] is how substitution events were identified relative to a reference sequence. Bhatt *et al.* [16] selected a sequence collected at an earlier date as the outgroup for each data set. Their analysis was not limited to viruses related by ladder-like trees. The underlying assumption is that mutations mapped to the branch from the outgroup to the in-group represent fixed differences driven by selection in an evolving population. In contrast, Kistler and Bedford [15] tracked a single trunk lineage over longer time periods for different virus species. To account for repeated substitutions at the same sites, they updated the reference sequence at successive intervals by taking the consensus of the preceding time interval. This implicitly assumes that the branch that roots the clade in the next time interval is determined by selection. Interestingly, Bhatt and colleagues also performed a similar study [58] that focused on proteins of human influenza A virus over three decades of evolution — taking the consensus of the earliest time point as the reference — and observed strong adaptive evolution in hemagglutinin where they did not using their previous approach [16]. A recent paper from Barrat-Charlaix and Neher [59] may shed light on this discrepancy. Using a simulation model, they found that heterogeneous and rapidly waning host immunity results in a decay of selective advantage that could explain the unpredictability of human influenza A virus evolution. As a result of this decay, the genetic makeup of the lineage that becomes the next trunk is predicted more by its relative frequency, *i.e.*, neutral evolution, than its original selective advantage. This suggests that some mutations associated with the trunk lineage may arise in the relative absence of selection. The majority of codon substitutions are non-synonymous. In addition, empirical evidence suggests that only a small number of sites in the surface-exposed proteins of influenza A virus are under positive selection by neutralizing antibodies [8, 60] (see also Figure 1B). Evaluating the impact of dynamic fitness effects on measuring adaptive evolution in this context will be an important area for further investigation.

### Limitations and future directions

Evolutionary fingerprints can provide a useful framework for comparing unrelated protein-coding genes with respect to how their diversities have been shaped by selection [19]. This method enables the investigator to compare site-specific rate estimates between unrelated genes through the common framework of their latent distributions. However, evolutionary fingerprinting is challenging to implement in practice. In this study, we have observed that fingerprints are sensitive to the amount of genetic variation in the data, which is quantified by tree length (also known as phylogenetic diversity). The length of the alignment has the same effect because it increases the sample size with respect to substitution events. Hence, both quantities determine the extent by which the likelihood can reshape the prior distribution. The original developers of the evolutionary fingerprinting method [19] also noted that resolving the distribution of site-specific rates was affected by sample size and genetic divergence. However, they used a different approach by fitting a general discrete distribution, in which *K* rate classes are each represented by three parameters: the class probability *p_k_* and class-specific *dN_k_* and *dS_k_* rates. These parameters were estimated from the data by maximum likelihood under the constraints 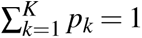 and 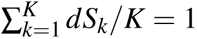. The issue of sample size was mitigated by approximating the posterior distribution around these point estimates. Our findings indicate that even adopting a fully Bayesian approach is not sufficient, and that normalizing the amount of information among datasets is necessary to account for large differences in sampled genetic variation among RNA viruses. In contrast, the study by Murrell *et al.* [21], on which we modeled our approach to evolutionary fingerprinting, used this method to characterize the coevolution between primate retroviruses and host restriction factors. Because their analysis comprised genes encoding restriction factors from the same set of mammalian genomes, it was not necessary to normalize the amount of genetic variation in their data.

A limitation of evolutionary fingerprinting is that it relies on a codon substitution model that is time homogeneous, *i.e.*, with constant *dN* and *dS* rates at each codon site. This means that selection pressures must be consistently maintained across multiple lineages over time to be detectable. However, site-specific selection pressures can change over time. This shift can be detected by branch-site or episodic selection models [61], although there is an inherent limit to their statistical power. Extending the evolutionary fingerprint to accommodate time-heterogeneous rates remains an open problem. Another limitation of evolutionary fingerprinting is that information about variation in *dN* and *dS* rates is collapsed into a bivariate distribution, which discards information about the relative locations of these codon sites in the gene. For example, positively-selected sites tend to cluster in the tertiary structures of proteins [62], and this rate variation is associated with solvent accessibility and residue contacts [63]. Incorporating positional information is difficult because of variation in sequence lengths among different genes. One possible solution might be to allow a gene to have multiple bivariate *dN*-*dS* distributions. These distributions could be mapped to codon sites by a hidden Markov model. We would then need to extend the Wasserstein distance to compare fixed lengths of subsequences of fingerprints between two gene alignments.

## Supplementary Figures

**Figure S1:**
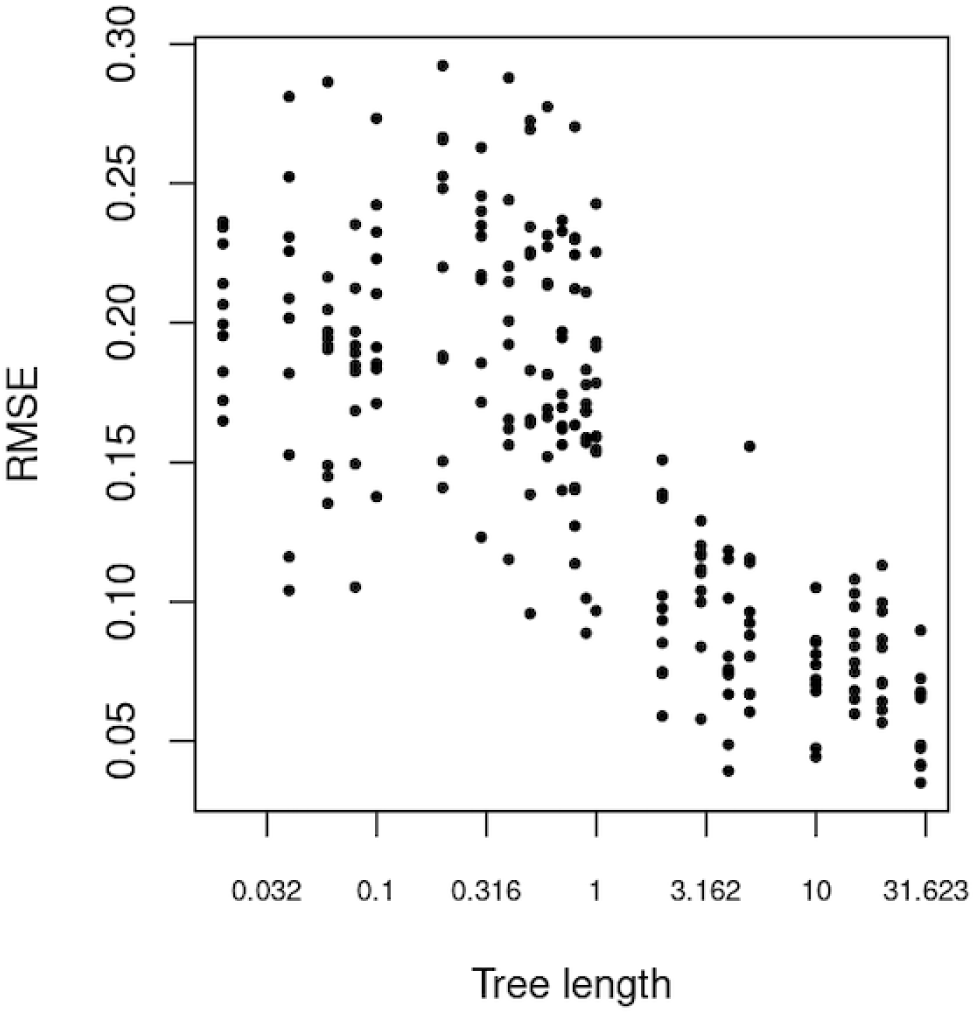
Accuracy in estimating site-specific dN/dS improves with tree length. Sequence alignments were simulated from a coalescent tree that was rescaled to different lengths, expressed in units of expected substitutions per codon site. These can be converted to expected substitutions per *nucleotide* site by dividing the value by 3. We calculated the root mean square error (RMSE) between the known dN/dS values and the estimated values across codon sites. Each point represents the RMSE for one of ten replicates per tree length, for 22 different lengths: 0.02, 0.04, 0.06, 0.08, 0.1, 0.2, 0.3, 0.4, 0.5, 0.6, 0.7, 0.8, 0.9, 1.0, 2.0, 3.0, 4.0, 5.0, 10.0, 15.0, 20.0, 30.0.

**Figure S2:**
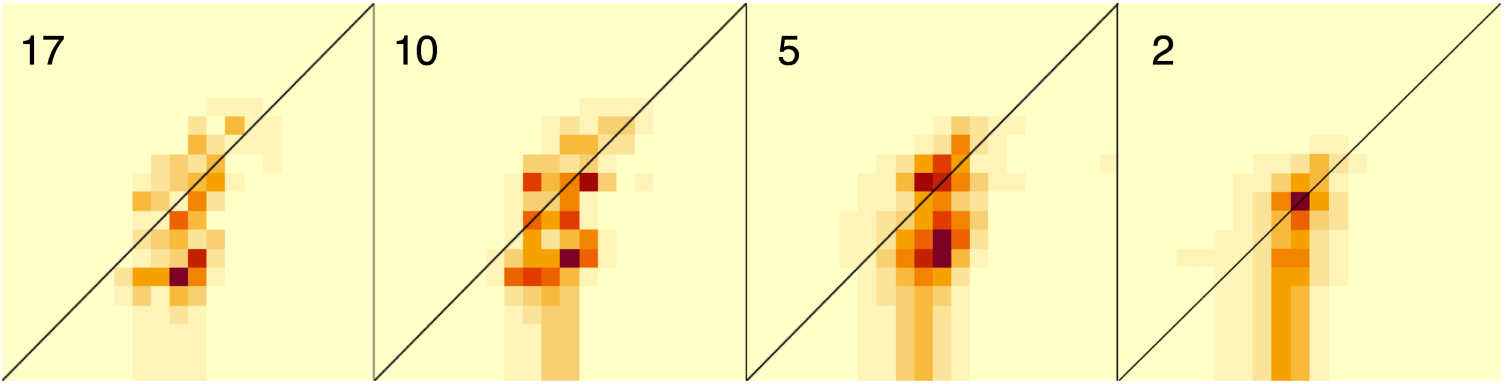
Effect of tree length of evolutionary fingerprints. Each fingerprint depicts the posterior probability distribution over a fixed grid of 20 × 20 *dS* (x-axis) and *dN* (y-axis) values. Darker cell shades correspond to higher posterior probabilities. These fingerprints were derived from progressively smaller numbers of HIV-1 *env* sequences, resulting in shorter tree lengths (as measured by the expected number of nucleotide substitutions, upper left).

**Figure S3:**
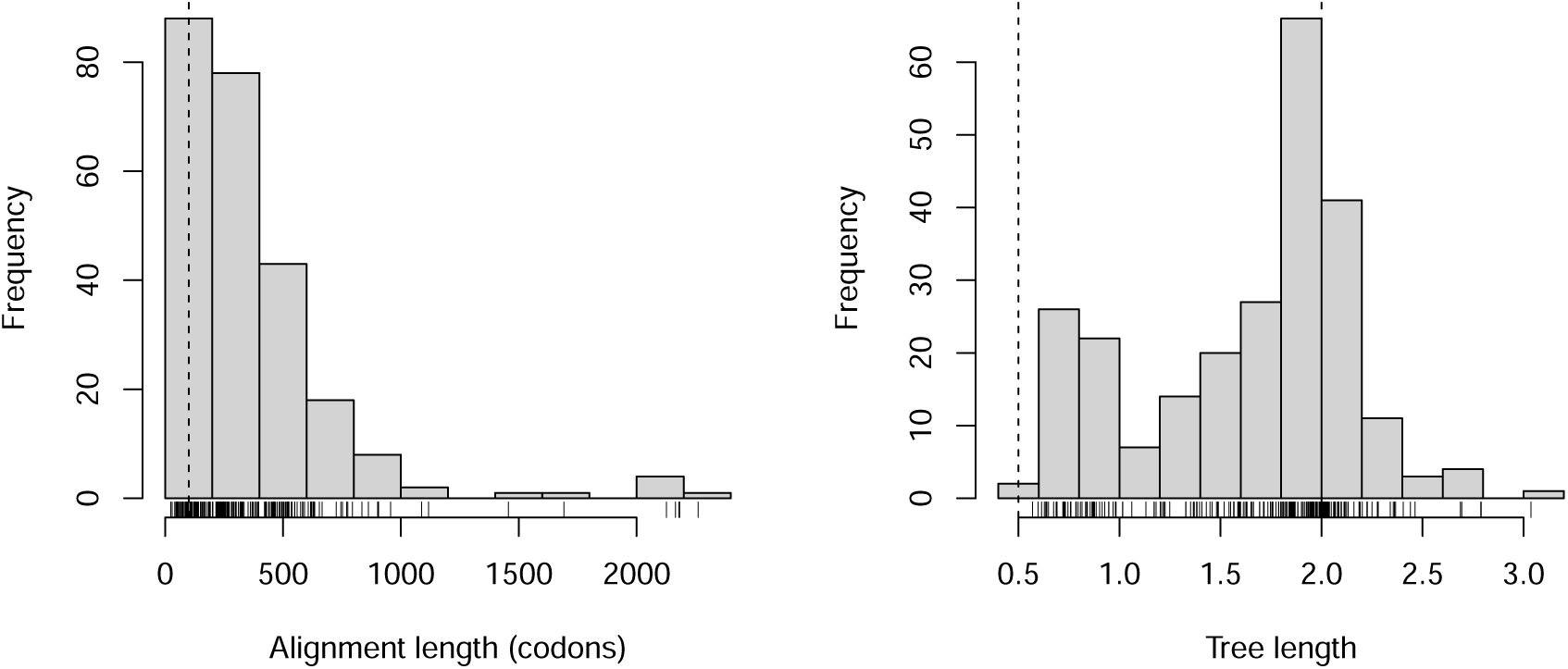
Distributions of alignment and tree lengths among *n* = 244 gene alignments. These histograms summarize the distributions of alignment length (in codons, left) and tree lengths (in expected number of substitutions per nucleotide site, right). Note that some tree lengths were slightly above our target of 2.0 substitutions per nucleotide site because removing the next branch would result in a tree length that was even further from this target.

**Figure S4:**
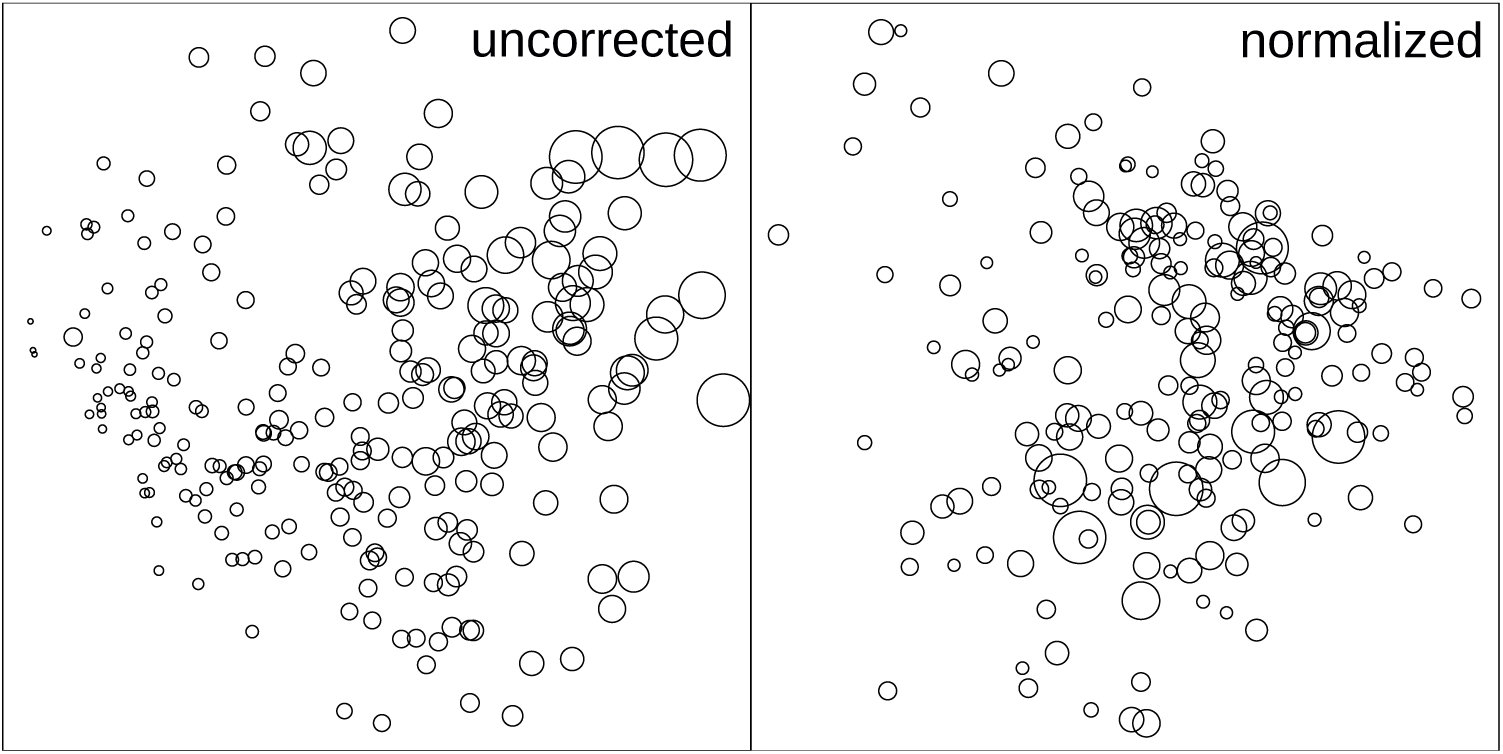
Effect of alignment length on evolutionary fingerprints. These plots depict the multidimensional scaling projection of the Wasserstein distance matrix. Each point represents a single gene alignment (uncorrected, *left*) or the centroid of 10 random samples of 100 codon sites from each alignment (normalized, *right*); alignments fewer than 100 codon in length were excluded from the latter. Point area is scaled to the number of codon sites in the original alignment.

**Figure S5:**
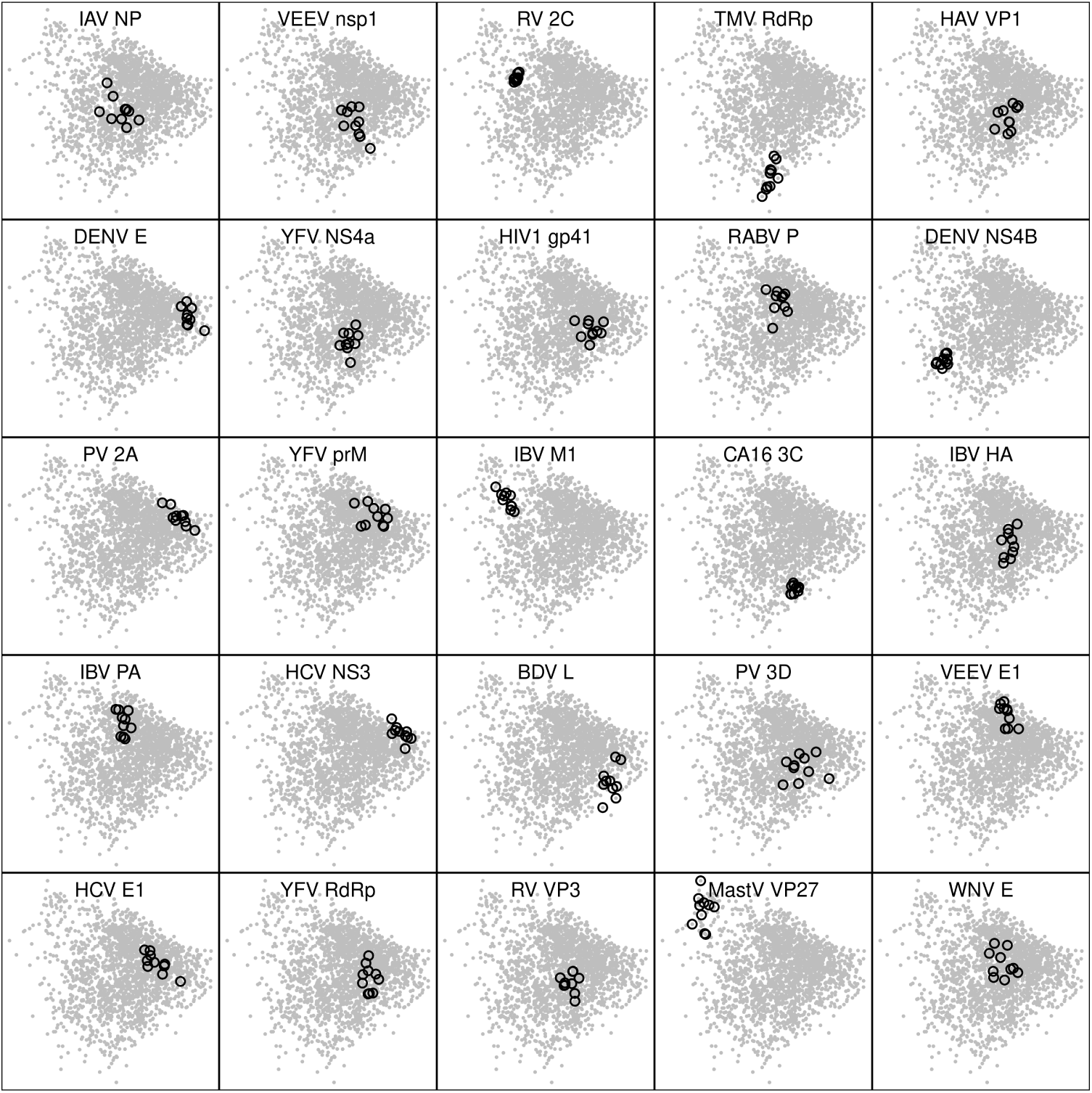
Clustering of fingerprints derived from random samples of alignments. Each plot is derived from the same multidimensional scaling projection of the Wasserstein distance matrix for samples of 100 codon sites from gene alignments. Points (open circles, black) corresponding to the 10 replicate samples from a given gene alignment are highlighted for a random selection of viruses and protein-coding genes (labels).

**Figure S6:**
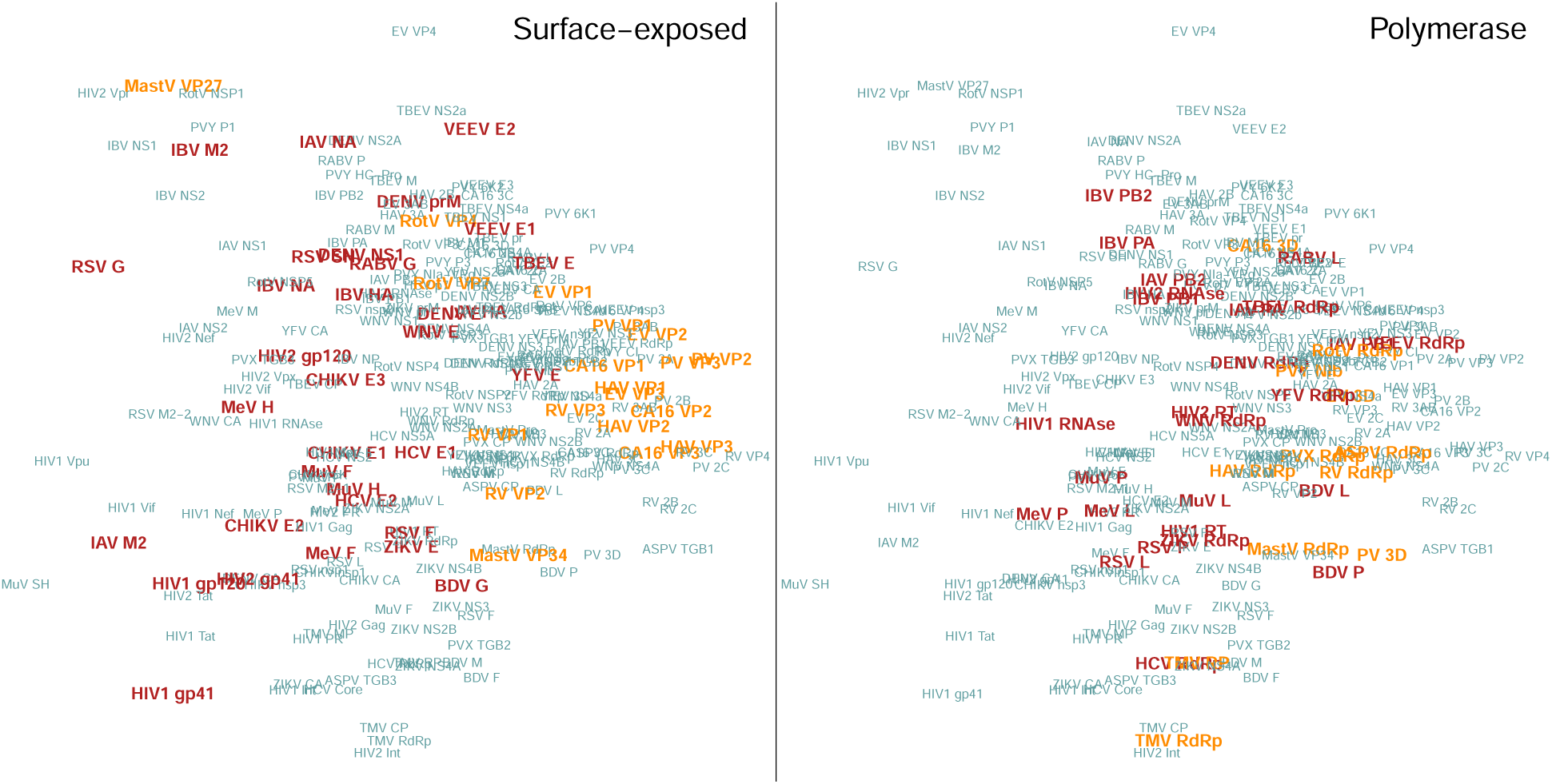
Reducing alignment length from 100 to 50 codons does not qualitatively affect results. Like Figure 3, these plots depict the multidimensional scaling projection of the Wasserstein distance matrix. Each point represents the centroid of 10 random samples of 50 codon sites from each alignment, labeled with abbreviations for virus and gene product. Labels are highlighted in bold for surface-exposed proteins (*left*) or proteins with polymerase activity (*right*), and with a darker shade (red) for enveloped viruses.

**Figure S7:**
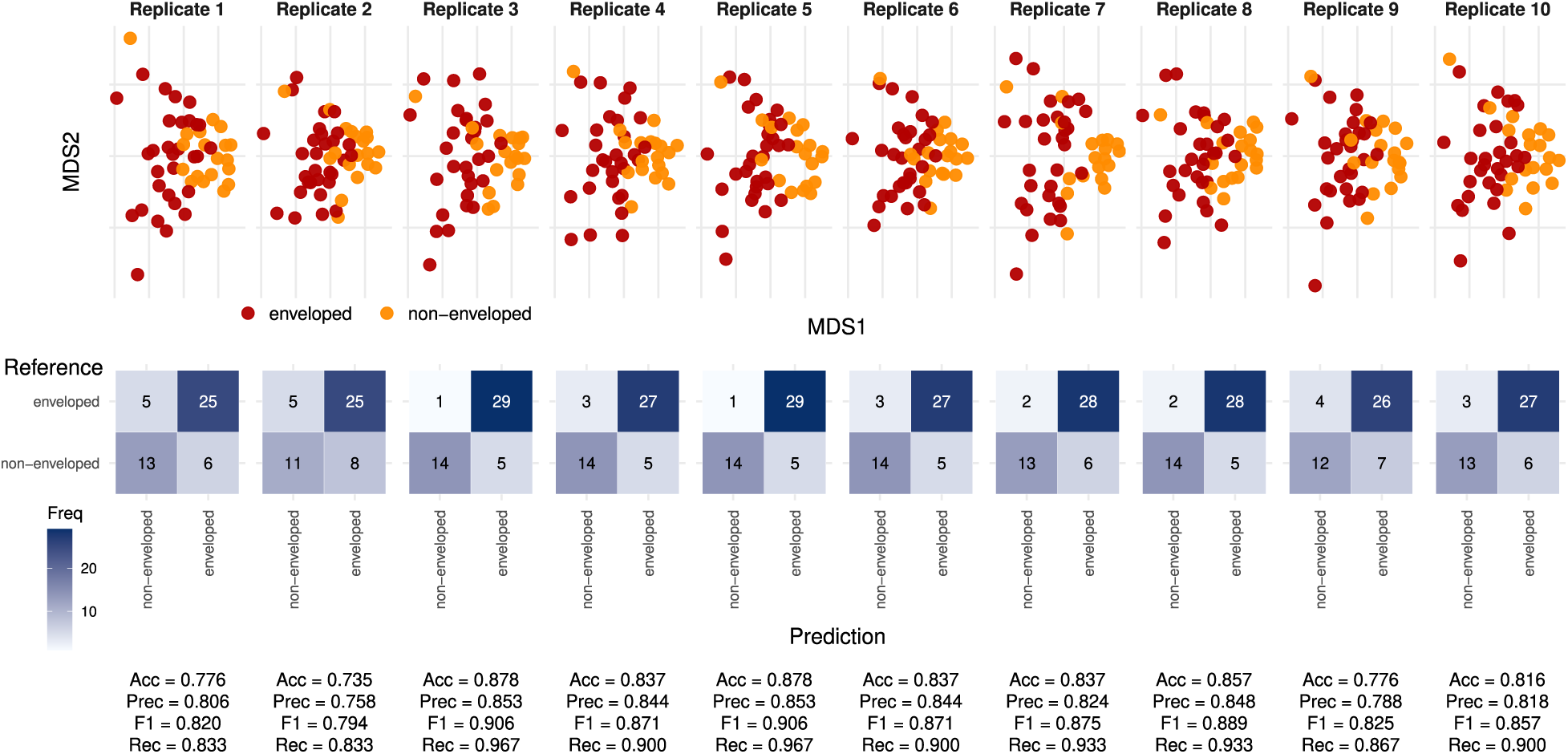
Multidimensional scaling (MDS) and confusion matrices for *k*-Nearest Neighbors (*k* = 5). The upper panel shows the two-dimensional MDS projections of protein distances for each of the ten independent replicates. Each point represents a protein classified as either *enveloped* (red) or *non-enveloped* (orange). The partial separation between clusters indicates that the distance matrix captures biologically meaningful relationships related to surface exposure and envelope association, although some overlap remains due to shared structural features. The lower panel displays the corresponding confusion matrices for each replicate, normalized by row percentages. Dark blue cells represent correctly classified instances, while lighter shades indicate misclassifications. Performance metrics (Accuracy, Recall, F1-score, and Precision) are reported below each matrix, showing consistent results across replicates (average Accuracy ≈ 84–85%). Together, these visualizations illustrate the reproducibility and robustness of the distance-based classification, confirming that the *k*NN model effectively distinguishes between enveloped and non-enveloped proteins.

**Figure S8:**
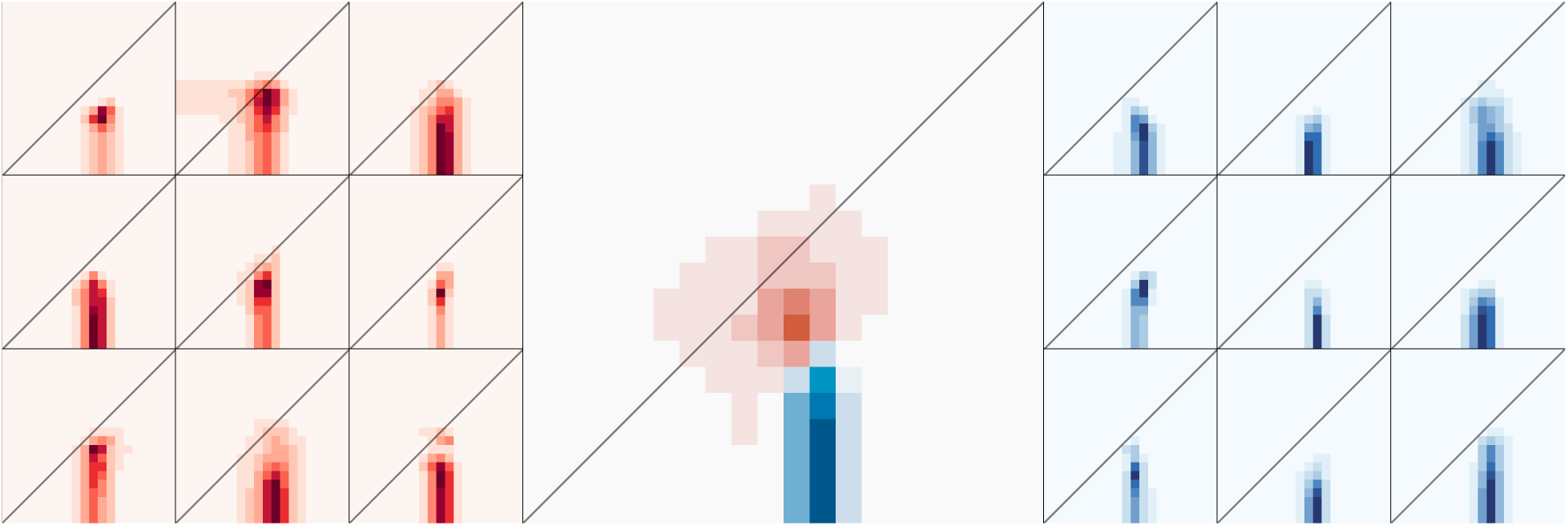
The difference in evolutionary fingerprints between enveloped and non-enveloped viruses. Evolutionary fingerprints inferred for surface-exposed proteins for a random selection of nine enveloped (left) or non-enveloped (right) viruses are displayed as heatmaps. A larger heatmap in the middle depicts the difference between evolutionary fingerprints averaged over *n* = 30 enveloped or *n* = 19 non-enveloped viruses. Blue-coloured cells indicate combinations of *dN* (y-axis) and *dS* (x-axis) rates with a higher posterior probability in non-enveloped viruses, and red cells have a higher probability in enveloped viruses.

**Figure S9:**
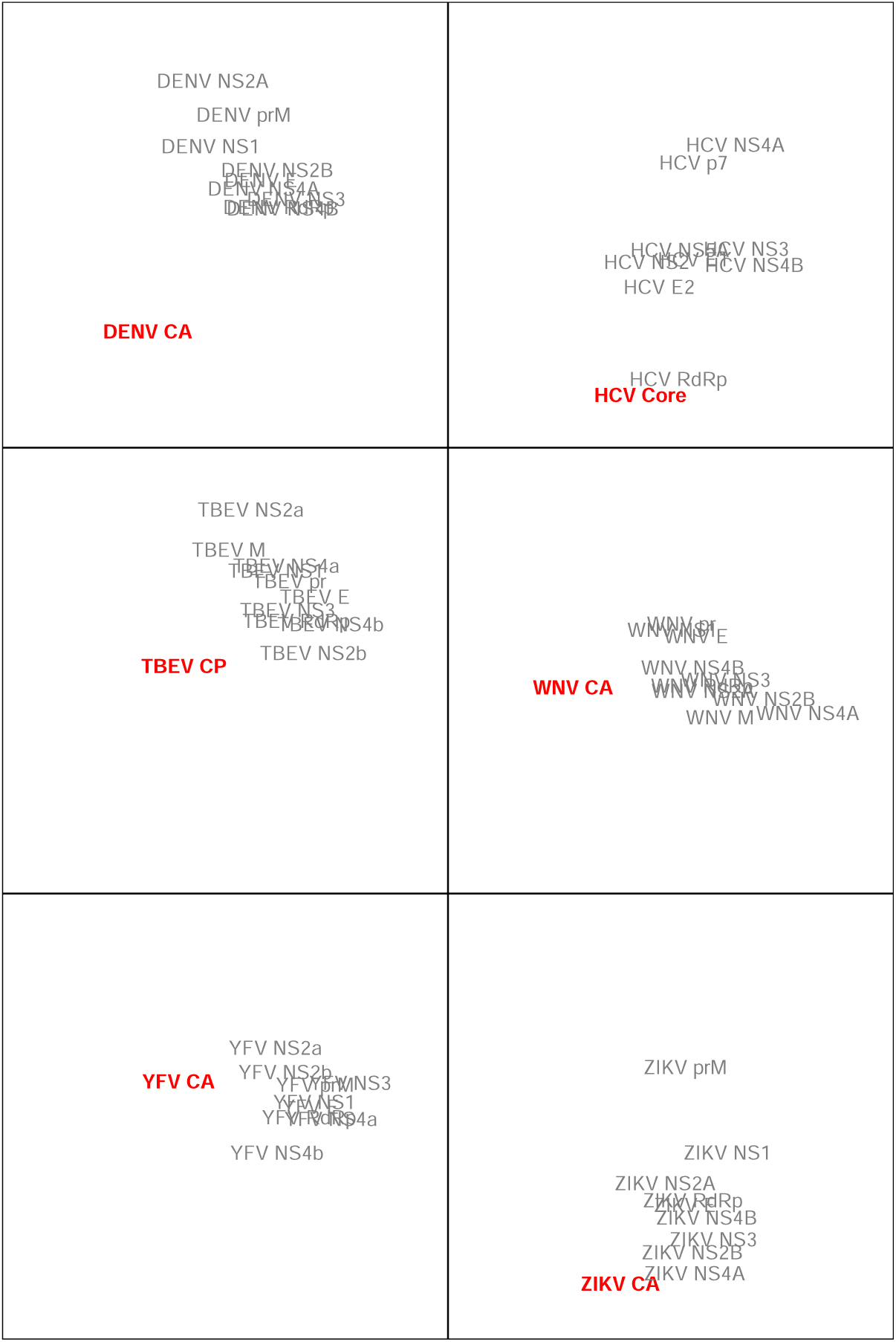
Multi-dimensional scaling projections of Wasserstein distance matrices for evolutionary fingerprints associated with flavivirus proteins. Each point represents the centroid of ten replicate samples of 50 codons from a protein-coding gene alignment. After removing overlapping regions, the hepatitis C virus (HCV) Core protein was too short to be included in our analysis of replicate *L* = 100 codon alignments.

## Supplementary Tables

**Table S1:**
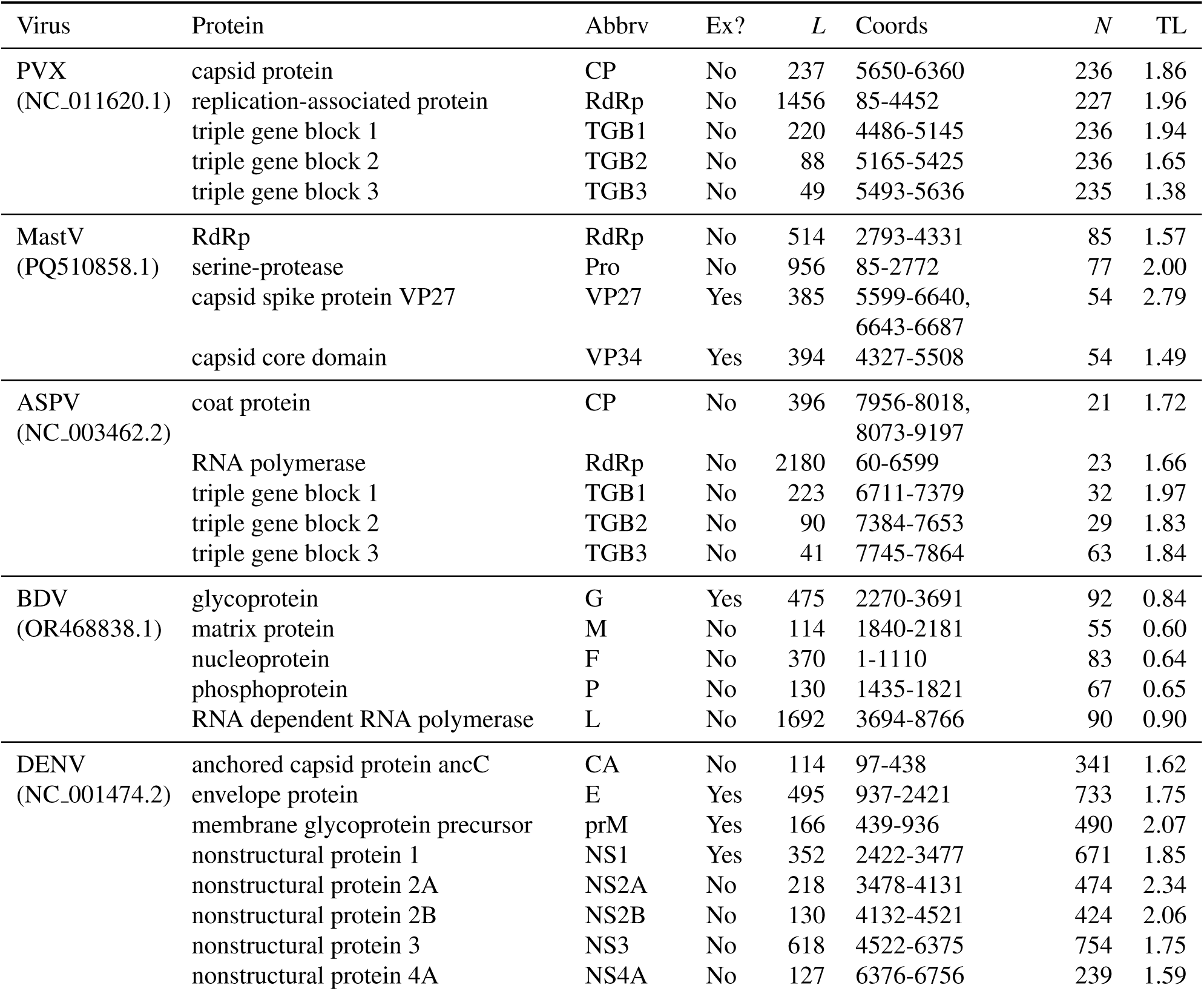
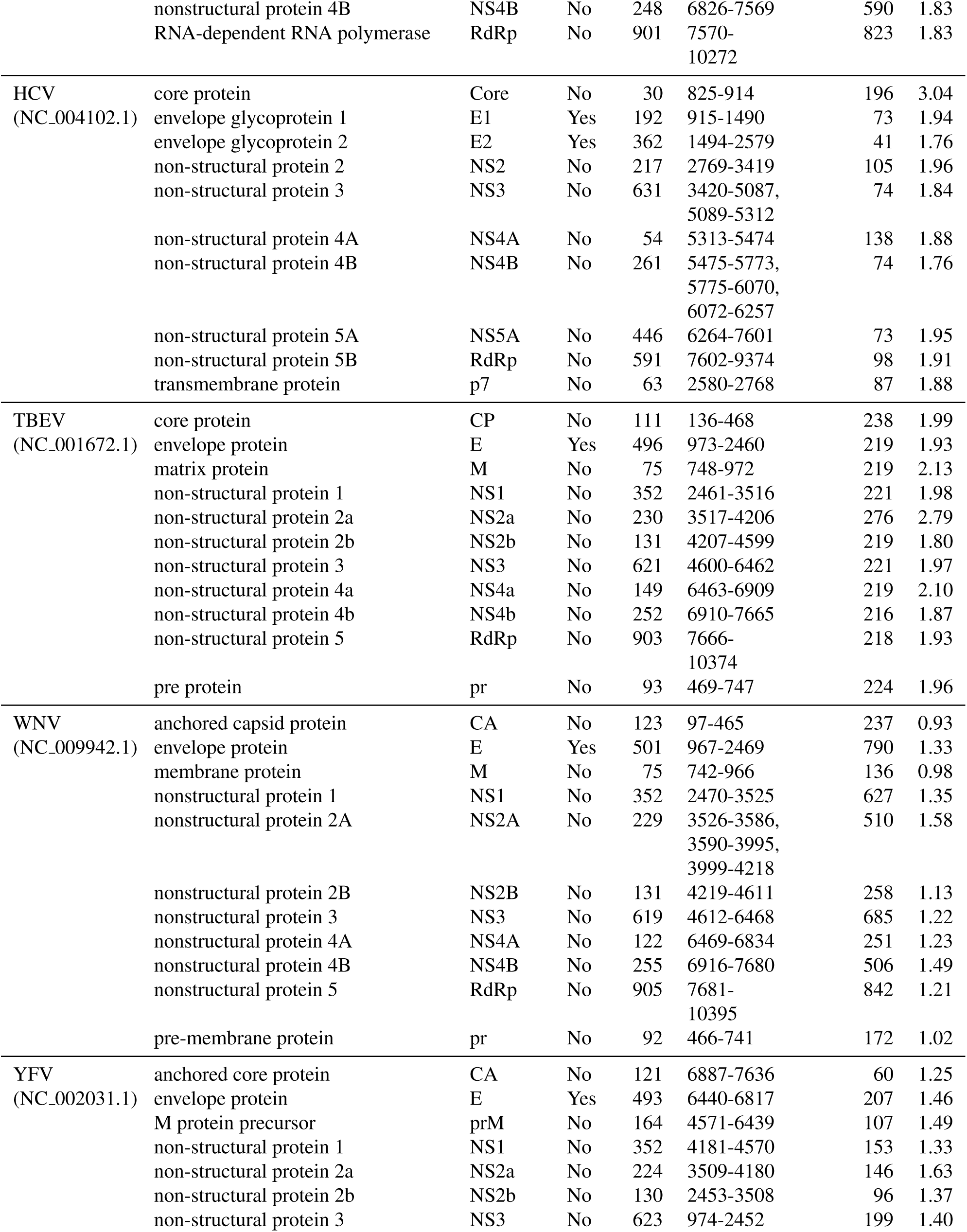
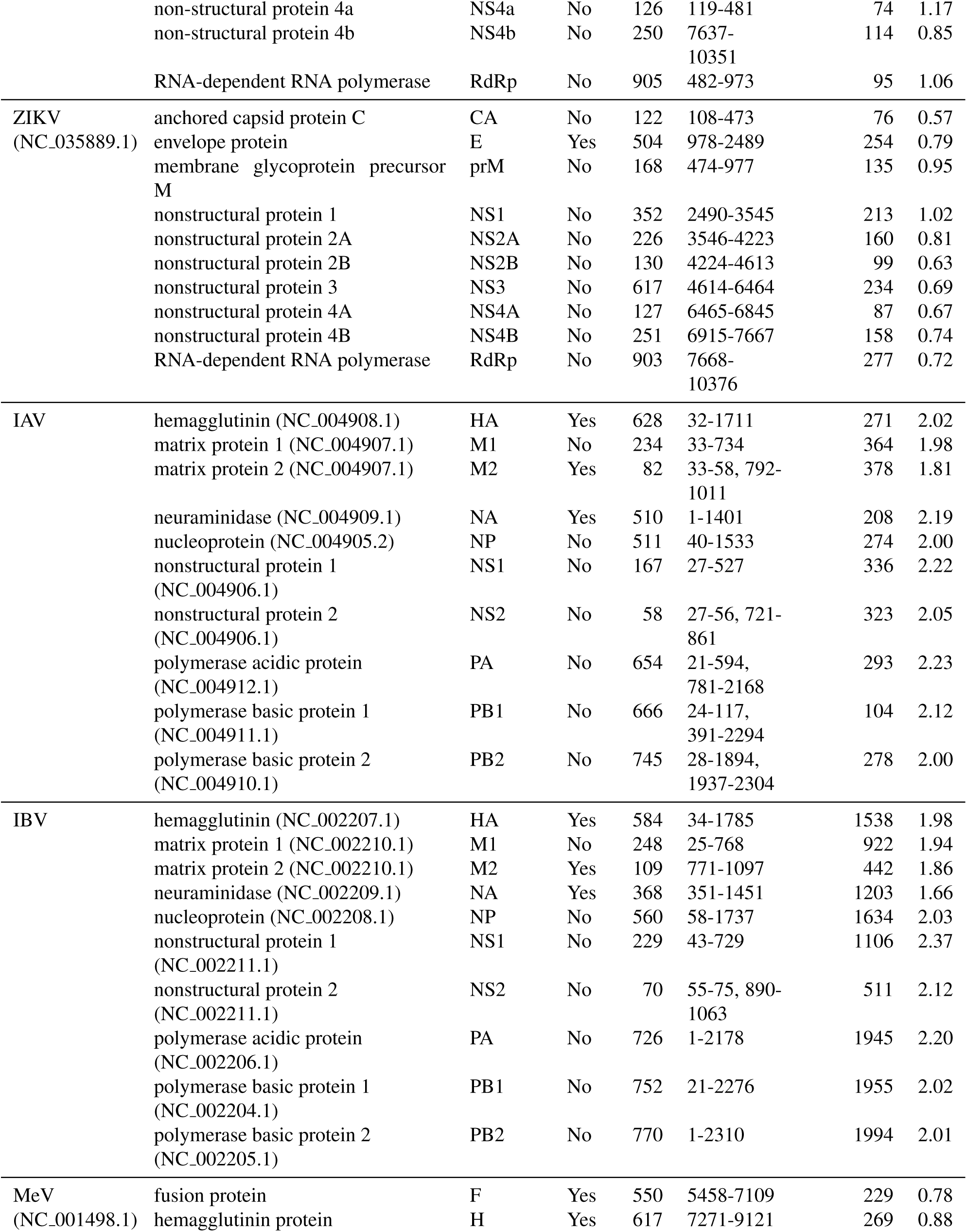
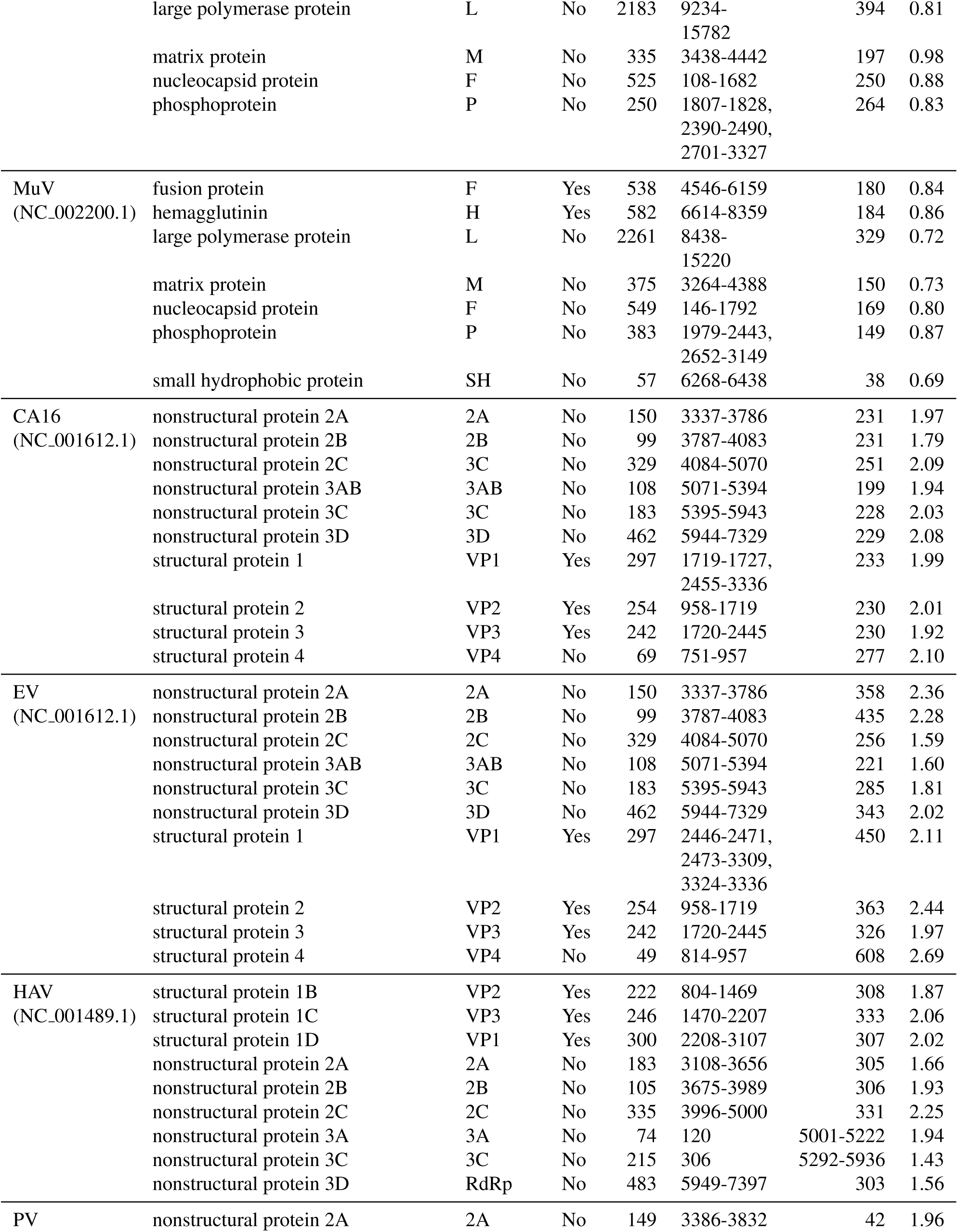
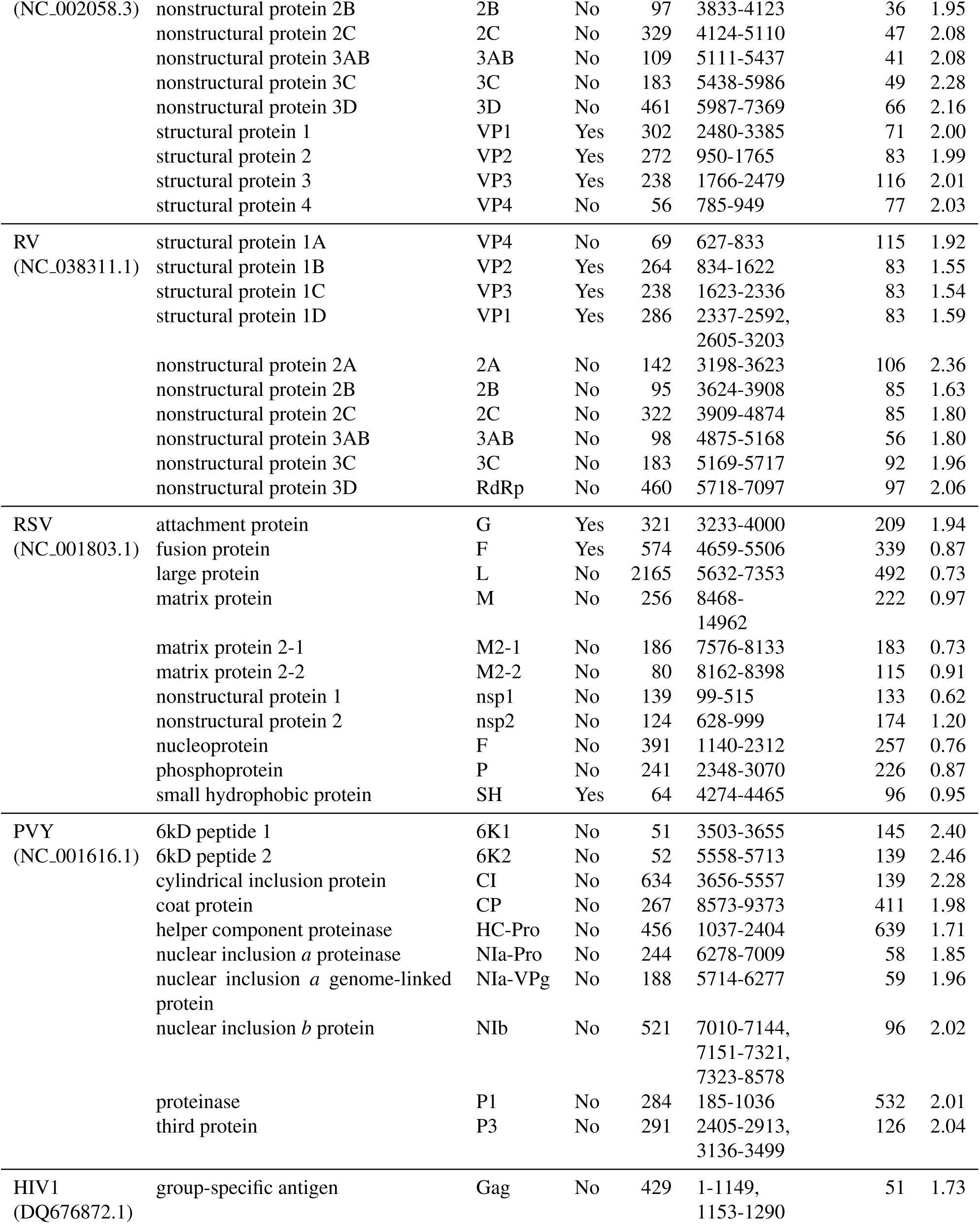
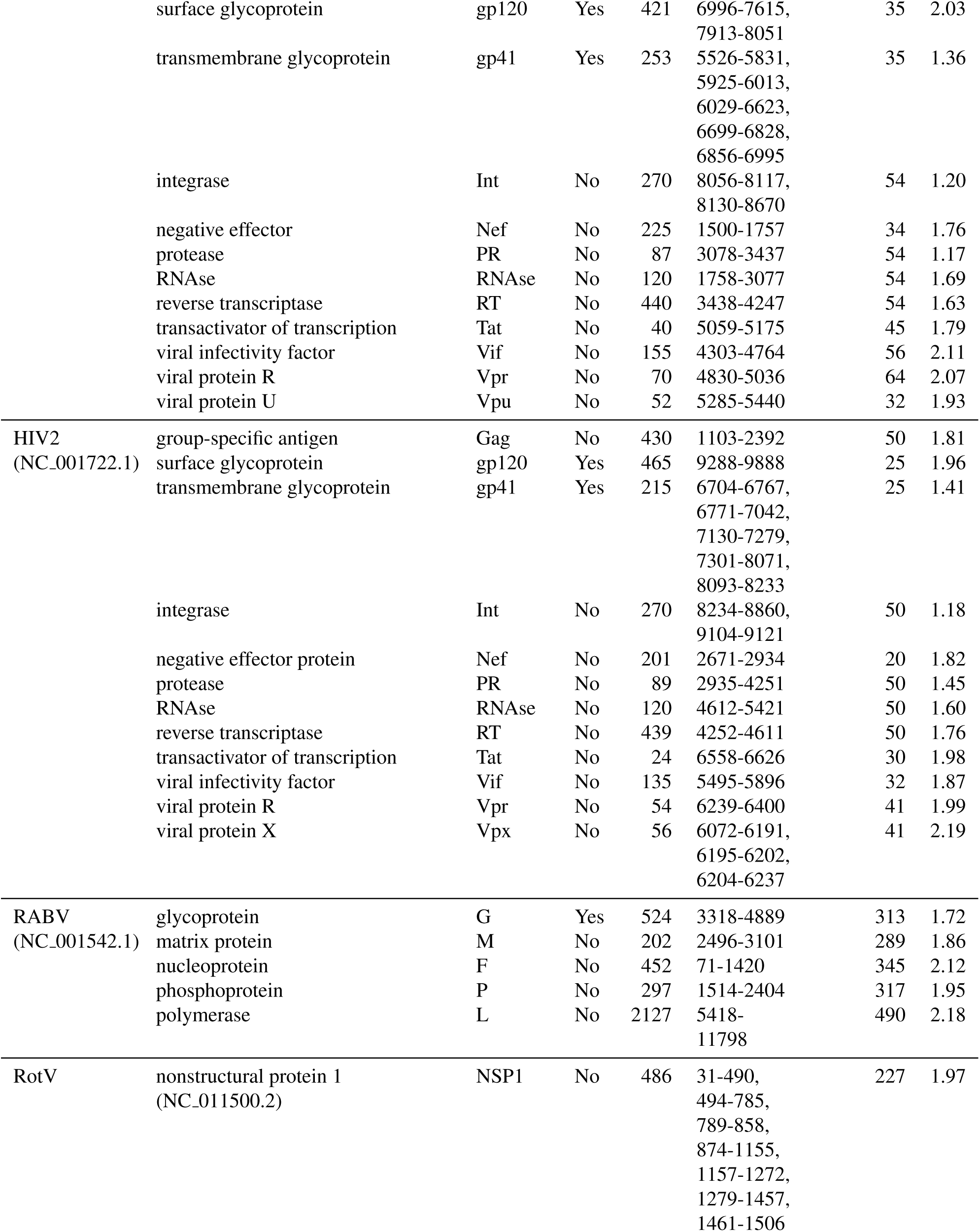
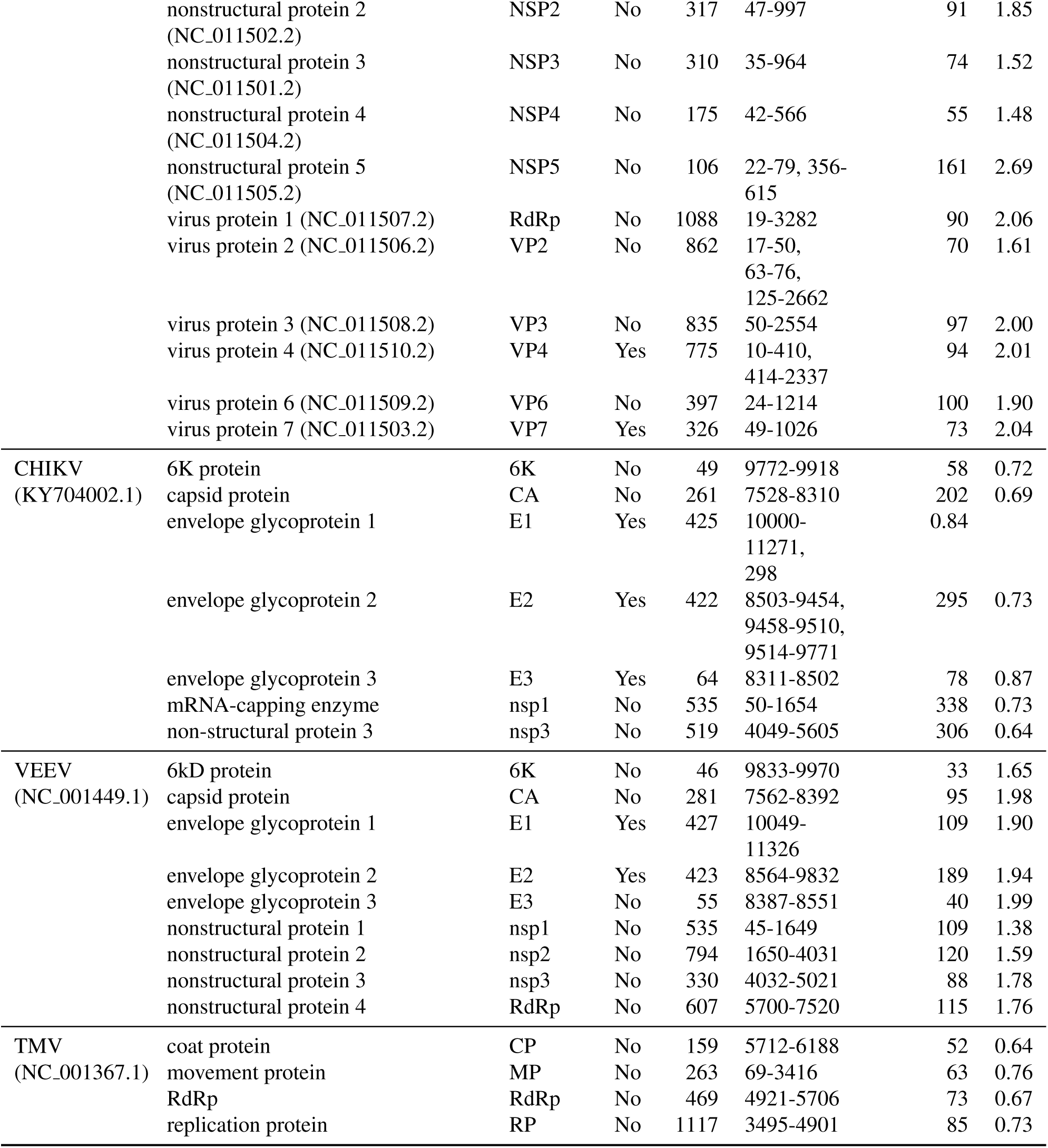
Summary of viral protein names and characteristics. *Virus* = abbreviated virus name (see Table 1). Followed by the Genbank accession of the reference genome used for determining gene coordinates; if the genome is segmented, then the accessions are provided alongside the respective gene products (*Protein*). *Abbrv* = abbreviation of protein name for figures. *Ex?* = is the protein classified as surface-exposed? *L* = the number of codon sites prior to normalizing alignment lengths by random sampling. *Coords* = nucleotide coordinates in reference genome, determined by pairwise alignment of the consensus sequence of the curated data set. Multiple ranges are given for products of spliced exons, to remove indels with respect to the reference, or when an overlapping open reading frame was removed from the alignment. *N* = the number of sequences after normalizing tree length. *TL* = total tree length (expected substitutions per nucleotide site) after normalization by pruning.

